# Estrogen-related receptor agonism reverses mitochondrial dysfunction and inflammation in the aging kidney

**DOI:** 10.1101/755801

**Authors:** Xiaoxin X. Wang, Komuraiah Myakala, Andrew E. Libby, Julia Panov, Suman Ranjit, Shogo Takahashi, Bryce A. Jones, Kanchan Bhasin, Yue Qi, Kristopher W. Krausz, Patricia M. Zerfas, Thomas J. Velenosi, Daxesh P. Patel, Parnaz Daneshpajouhnejad, Avi Titievsky, Vadim Sharov, Boris Ostretsov, Cyrielle Billon, Arindam Chatterjee, John K. Walker, Jeffrey B. Kopp, Avi Z. Rosenberg, Frank J. Gonzalez, Udayan Guha, Leonid Brodsky, Thomas P. Burris, Moshe Levi

**Affiliations:** Department of Biochemistry and Molecular & Cellular Biology, University of Haifa, Mount Carmel, Haifa, 31905, Israel; Tauber Bioinformatics Research Center, University of Haifa, Mount Carmel, Haifa, 31905, Israel; Sagol Department of Neurobiology, University of Haifa, Mount Carmel, Haifa, 31905, Israel; Laboratory of Metabolism, Center for Cancer Research, National Cancer Institute, National Institutes of Health, Bethesda, MD 20892, USA; Department of Pharmacology and Physiology, Georgetown University, Washington DC 20057, USA; Thoracic and GI Malignancies Branch, National Cancer Institute, National Institutes of Health, Bethesda, MD 20814, USA; Office of Research Services, Office of the Director, National Institutes of Health, Bethesda, MD 20892, USA; Department of Pathology, Johns Hopkins University School of Medicine, Renal Pathology Service, Baltimore, MD 21287, USA; Center for Clinical Pharmacology, Washington University School of Medicine and St. Louis College of Pharmacy, St. Louis, MO 63110, USA; Department of Pharmacology & Physiology, Saint Louis University School of Medicine, St. Louis, MO 63104, USA; Kidney Diseases Section, National Institute of Diabetes and Digestive and Kidney Diseases, National Institutes of Health, Bethesda, MD 20814, USA

## Abstract

**Background:** A gradual decline in renal function occurs even in healthy aging individuals. In addition to aging *per se*, concurrent metabolic syndrome and hypertension, which are common in the aging population, can induce mitochondrial dysfunction and inflammation, which collectively contribute to age-related kidney dysfunction and disease. Here we studied the role of the nuclear hormone receptors, the estrogen-related receptors (ERRs) in regulation of age-related mitochondrial dysfunction and inflammation. ERRs were decreased in aging human and mouse kidneys and were preserved in aging mice with lifelong caloric restriction (CR).

**Methods:** A pan-ERR agonist was used to treat 21-month-old mice for 8-weeks. In addition, 21-month-old mice were treated with a STING inhibitor for 3 weeks.

**Results:** Remarkably, only an 8-week treatment with a pan-ERR agonist reversed the age-related increases in albuminuria, podocyte loss, mitochondrial dysfunction and inflammatory cytokines, including the cGAS-STING and STAT3 signaling pathways. A 3-week treatment of 21-month-old mice with a STING inhibitor reversed the increases in inflammatory cytokines and the senescence marker p21 but also unexpectedly reversed the age-related decreases in PGC-1α, ERRα, mitochondrial complexes and MCAD expression.

**Conclusions:** Our studies identified ERRs as important modulators of age-related mitochondrial dysfunction and inflammation. These findings highlight novel druggable pathways that can be further evaluated to prevent progression of age-related kidney disease.

**Significance Statement:** There is an increasing need for prevention and treatment strategies for age-related kidney disease. The hallmarks of aging kidneys are decreased mitochondrial function and increased inflammation. The expression of the nuclear hormone receptors estrogen-related receptors (ERRs) are decreased in aging human and mouse kidneys. This paper investigates the role of ERRs in the aging kidney. Treatment of aging mice with a pan-ERR agonist reversed the age-related increases in albuminuria and podocyte loss, mitochondrial dysfunction and inflammatory cytokines, including the cGAS-STING signaling pathways. Treatment of aging mice with a STING inhibitor decreased inflammation and increased mitochondrial gene expression. These findings identify ERRs as important modulators of age-related mitochondrial dysfunction and inflammation.

## INTRODUCTION

The fastest growing segment of the US population with impaired kidney function is the 65 and older age group. This population is expected to double in the next 20 years, while the number worldwide is expected to triple from 743 million in 2009 to 2 billion in 2050. This will result in a marked increase in the elderly population with chronic kidney disease. This increase may be further amplified by other age-related co-morbidities including metabolic syndrome and hypertension that accelerate age-related decline in renal function ^1^. Thus, there is a growing need for prevention and treatment strategies for age-related kidney disease.

A gradual age-related decline in renal function occurs even in healthy aging individuals ^2^. Progressive glomerular, vascular and tubulointerstitial sclerosis is evident on renal tissue examination of healthy kidney donors with increasing age ^3^. In addition to aging *per se*, metabolic syndrome and hypertension can induce mitochondrial dysfunction and inflammation, as well as endoplasmic reticulum stress, oxidative stress, altered lipid metabolism, and elevation of profibrotic growth factors in the kidney, which collectively contribute to age-related kidney disease ^2^.

There is variation in the rate of decline in renal function as a function of sex, race, and burden of co-morbidities ^4–6^. Interestingly, examination of processes leading to sclerosis suggests a role for possible modifiable systemic metabolic and hormonal factors that can ameliorate the rate of sclerosis. With the population of older individuals increasing, identifying preventable or treatable aspects of age-related nephropathy becomes of critical importance.

There is increasing evidence that mitochondrial biogenesis, mitochondrial function, mitochondrial unfolded protein response (UPR^mt^), mitochondrial dynamics, and mitophagy are impaired in aging, and these alterations contribute to the pathogenesis of the age-related diseases ^7–9^. In this regard current studies are concentrated on modulating these molecular mechanisms to improve mitochondrial function.

Caloric restriction (CR) plays a prominent role in preventing age-related complications. We have previously shown that CR prevents the age-related decline in renal function and renal lipid accumulation via, at least in part, inhibition of the increased activity of the sterol regulatory element binding proteins (SREBPs) ^10, 11^. CR is also an important modulator of mitochondrial function. We have demonstrated that CR prevents age-related mitochondrial dysfunction in the kidney by increasing mitochondrial/nuclear DNA ratio, mitochondrial complex activity including fatty acid β-oxidation and expression of the mitochondrial transcription factor NRF1, the protein kinase AMPK, the deacetylases sirtuin 1 and 3, and mitochondrial isocitrate dehydrogenase (IDH2) expression ^12^. In addition, CR also prevents the age-related decrease in mitochondrial number in the renal tubules. CR increases expression of the bile acid regulated nuclear receptor farnesoid X receptor (FXR) and the Takeda G protein coupled receptor (TGR5). Treatment of 22-month-old *ad lib* (AL)-fed mice for 2 months with the dual FXR-TGR5 agonist INT-767 reversed most of the age-related impairments in mitochondrial function and the progression of renal disease ^12^. Importantly INT-767 and CR each increased expression of PGC-1α, ERRα, and ERRγ, which are important regulators of mitochondrial biogenesis and function.

The estrogen-related receptors (ERRs) ERRα (NR3B1, ESSRA gene), ERRβ (NR3B2, ESRRB gene), and ERRγ (NR3B3, ESRRG gene) are members of the nuclear receptor superfamily. While one report suggests a role for cholesterol ^13^, there are no confirmed endogenous ligands for these orphan receptors. Importantly the ERRs do not bind natural estrogens, and they do not directly participate in classic estrogen signaling pathways or biological processes ^14^. ERRα and ERRγ are strongly activated by their coactivators PGC-1α and PGC-1β ^15^. In contrast, RIP140 and NCoR1 are important corepressors of ERR activity ^16, 17^. ERRs are subject to post-translational modifications including phosphorylation, sumoylation, and acetylation that modulate their DNA binding and recruitment of coactivators ^18, 19^.

ERRα and ERRγ regulate the transcription of genes involved in mitochondrial biogenesis, oxidative phosphorylation, tricarboxylic acid (TCA) cycle, fatty acid oxidation and glucose metabolism ^14^. However, in addition to overlapping gene activation there is also ample evidence that ERRα and ERRγ have differential and opposing effects, which can be due to interactions with corepressors, coactivators, posttranslational modification, or differential cell expression ^14^. Opposing effects for ERRα and ERRγ are seen in breast cancer ^20^, regulation of gluconeogenesis in the liver ^20, 21^, skeletal muscle function ^20, 21^, macrophage function ^21, 22^, and regulation of lactate dehydrogenase A (LDHA) related to anaerobic glycolysis ^14^.

ERRα and ERRγ are highly expressed in the mouse and human kidney ^23, 24^. However, the roles of ERRα and ERRγ in modulating age-related impairment of mitochondrial function and age-related inflammation (inflammaging) are not known. We undertook our current studies to determine whether a pan-ERR agonist, activating ERRα, ERRβ, and ERRγ, would improve mitochondrial function and inflammation in the aging mouse kidney.

## METHODS

### Mice

Studies were performed in 4-month-old and 21-month-old male C57BL/6 male mice obtained from the NIA aging rodent colony. Mice received 3% DMSO vehicle or the pan-ERR agonist SLU-PP-332 (Dr. Thomas Burris, Washington University) at a dose of 25 mg/kg body weight/day, administered intraperitoneally. Mice were dosed for 8 weeks, following which they underwent euthanasia and the kidneys were harvested and processed for a) histology, b) transmission electron microscopy, c) isolation of nuclei, d) isolation of mitochondria, and e) biochemical studies detailed below. Another cohort of male C57BL/6 mice with the same age were received the same vehicle as above or the STING inhibitor C-176 (Focus Biomolecules, Plymouth Meeting, PA) at a dose of 1mg/kg body weight/day for 3 weeks via intraperitoneal injection.

### Immunohistochemistry

Formalin-fixed paraffin-embedded tissue sections were subjected to antigen retrieval with EDTA buffer in high pressure heated water bath and staining was performed using either mouse monoclonal ERRα (1:2500, Abcam, Cambridge, MA) or ERRγ (1:400, Abcam) antibodies for 90 minutes or pyruvate dehydrogenase (PDH) E2/E3bp (1:1000, Abcam) antibody for 45 minutes. UnoVue HRP secondary antibody detection reagent (Diagnostic BioSystems, Pleasanton, CA) was applied followed by DAB chromogen. Imaging was done with Nanozoomer (Hamamatsu Photonics, Japan).

### Cell culture

Primary human proximal tubule epithelial cells (cat#: PCS-400-010) were purchased from ATCC (Manassas, VA). Cells were cultured in Renal Epithelial Cell Basal Medium (ATCC cat#: PCS-400-030) supplemented with Renal Epithelial Cell Growth Kit (ATCC cat#: PCS-4000-040) at 37°C in 5% CO2. Cells were cultured to 70-80% confluence and then treated with vehicle (4mM HCl with 0.1% bovine serum albumin) or 10 ng/mL TGF-β1 (Cat#: 7754-BH-005, R&D Systems, Minneapolis, MN) or 10 ng/mL TNF-α (Cat#210-TA-020, R&D Systems) for 24 hours. Cells were then harvested and analyzed for gene expression.

### Transmission Electron Microscopy

One mm^3^ cortex kidney tissues were fixed for 48 hrs. at 4°C in 2.5% glutaraldehyde and 1% paraformaldehyde in 0.1M cacodylate buffer (pH 7.4) and washed with cacodylate buffer three times. The tissues were fixed with 1% OsO_4_ for two hours, washed again with 0.1 M cacodylate buffer three times, washed with water and placed in 1% uranyl acetate for one hour. The tissues were subsequently serially dehydrated in ethanol and propylene oxide and embedded in EMBed 812 resin (Electron Microscopy Sciences, Hatfield, PA). Thin sections, approximately 80 nm, were obtained by utilizing the Leica ultracut-UCT ultramicrotome (Leica, Deerfield, IL) and placed onto 300 mesh copper grids and stained with saturated uranyl acetate in 50% methanol and then with lead citrate. The grids were viewed with a JEM-1200EXII electron microscope (JEOL Ltd, Tokyo, Japan) at 80kV and images were recorded on the XR611M, mid mounted, 10.5M pixel, CCD camera (Advanced Microscopy Techniques Corp, Danvers, MA). Mitochondrial morphology was assessed with ImageJ software by manually tracing all mitochondria that were completely within the field of view in 6 random images from each mouse (n = 3-4 each group). The area, perimeter, and minimum Feret diameter of each mitochondrion were measured.

### Autofluorescence FLIM

autofluorescence FLIM signals were acquired using an Olympus FVMPERS microscope (Waltham. MA) equipped with the DIVER (Deep Imaging via Enhanced-Photon Recovery) detector which was developed at the Laboratory for Fluorescence Dynamics (LFD), University of California at Irvine, CA, and FLIMBox (ISS, Champaign, IL) for phasor lifetime acquisition, as detailed in **Supplementary Methods**.

### RNA extraction and real-time quantitative PCR

Total RNA was isolated from the kidneys using Qiagen RNeasy mini kit (Valencia, CA), and cDNA was synthesized using reverse transcript reagents from Thermo Fisher Scientific (Waltham, MA). Quantitative real-time PCR was performed as previously described ^25^, and expression levels of target genes were normalized to 18S level. Primer sequences are listed in Supplementary Table 1.

### Bulk RNA-seq

Approximately 300-500 ng of kidneys RNA were used to generate barcoded RNA libraries using Ion AmpliSeq Transcriptome Mouse Gene Expression Panel, Chef-Ready Kit. Library quantification was performed using the Ion Library Quantitation Kit (Thermo Fisher Scientific). Sequencing was done on an Ion Proton with signal processing and base calling using Ion Torrent Suite (Thermo Fisher Scientific). Raw reads were mapped to Ampliseq supported mm10 transcriptome. Normalized read counts per million were generated using the RNA-seq Analysis plugin (Ion Torrent Community, Thermo Fisher Scientific). Gene expression table was converted to LN scale and quantile normalized. Genes with expression levels less than 4 across all samples were filtered out. Selection of the differentially expressed genes was performed utilizing student t-test. The significance of differentiation was defined as Student t-test statistic p-value < 0.05. Enrichment analysis of GO biological processes and pathways was prepared with DAVID ^26^ and with PANTHER online tool^27^. Heat map visualization was performed with Heatmapper tool ^28^.

### Single nuclei RNA-seq

Mouse kidney single nuclei were isolated ^29, 30^ and counted using the EVE Automated Cell Counter (NanoEnTek, VWR, Radnor, PA). The resulting mixture was provided to the Genomics and Epigenomics Shared Resource (GESR) at Georgetown University, and further processed by the Chromium Controller (10X Genomics, Pleasanton, CA) using Single Cell 3’ GEM Kit v3, Single Cell 3’ Library Kit v3, i7 multiplex kit, Single Cell 3’ Gel Bead Kit v3 (10X Genomics) according to the manufacturer’s instructions with modifications for single nuclei. Libraries were sequenced on the Illumina Novaseq S4 System (Illumina, San Diego, CA) to an average depth of >300 M reads PF per sample. Sequencing data were processed by Cellranger pipeline. Briefly, reads were aligned against the mouse mm10 genome reference with STAR algorithm. Barcodes and UMIs were filtered and corrected. Only confidently mapped, non-PCR duplicates with valid barcodes and UMIs were used to generate a gene-barcode matrix for further analysis. Expression matrix was further investigated by the Loupe Cell Browser.

### Proteomics

200 microgram of kidney tissue were homogenized and lysed by 8M urea in 20mM HEPES (pH=8.0) buffer with protease and phosphatase inhibitors using Tissue Lyser II (QIAGEN). Samples were reduced and alkylated followed by MS-grade trypsin digestion. The resulting tryptic peptides were labeled with 11 plex tandem mass tag (TMT). After quench, the tagged peptides were combined and fractionated with basic-pH reverse-phase high-performance liquid chromatography, collected in a 96-well plate and combined for a total of 12 fractions prior to desalting and subsequent liquid chromatography−tandem mass spectrometry (LC−MS/MS) processing on a Orbitrap Q-Exactive HF (Thermo Fisher Scientific) mass spectrometer interfaced with an Ultimate 3000 nanoflow LC system ^31^. Each fraction was separated on a reverse phase C_18_ nano-column (25cm × 75μ m, 2μ m particles) with a linear gradient 4∼45% solvent B (0.1% TFA in Acetonitrile). Data dependent mode was applied to analyze the top 15 most abundant peaks in one acquisition cycle.

MS raw files were mapped against Uniprot mouse database (version 20170207) using the MaxQuant software package (version 1.5.3.30) with the Andromeda search engine^32^. Corrected intensities of the reporter ions from TMT labels were obtained from the MaxQuant search. The normalized relative abundance of a protein in each sample was calculated as the ratio of a protein abundance in a sample to abundance of the same protein in a pool (channel 11, pooled from 20 samples) of the corresponding batch. After performing principle component analysis (PCA) on the protein relative abundance matrix we noticed that the samples were still separated by batches on the PC1-PC2 plane. Thus, we further corrected this batch effect by applying the Experimental Bayes (EB) batch correction method ^33^. After batch correction, PCA revealed clear separation of samples into biological groups on the PC1-PC2 plane (**Supplementary Figure 1**). All further analyses were performed on batch corrected protein abundance matrix. Selection of the differentially abundant proteins was done utilizing Wilks theorem based likelihood ratio test ^34^ and Student t-test statistic. The significance of differentiation was defined as likelihood ratio statistic p-value <0.001 and Student t-test statistic p-value < 0.05.

### Mitochondrial isolation

Kidney mitochondria were isolated using the kit (MITOISO1) from Sigma (St. Louis, MO) according to manufacturer instructions.

### Mitochondrial Biogenesis

We measured mitochondrial (*Cytb*) and nuclear (*H19*) DNA by RT-QPCR.

### Mitochondrial Respiration

We measured basal respiration, ATP turnover, proton leak, maximal respiration and spare respiratory capacity using the *Seahorse* XF96 Analyzer on equally loaded freshly isolated kidney mitochondria. We also measured mitochondrial complex I, II, III, IV, and V protein abundance by Native Blue Gel Electrophoresis (Thermo Fisher Scientific) with equally loaded mitochondrial fractions.

### Multi-omics data analysis and integration bioinformatics methods

Taking advantage of the multi-omics (transcriptomics and proteomics) measurements reflecting the same biological processes in the samples, we performed Two-way Orthogonal Partial Least Square (O2PLS) integration ^35^ to identify networks of associated genes and proteins. This analysis identified one orthogonal component (V2) that reflected the expected separation of samples into three distinct groups: young together with young pan-ERR agonist treated mice, old mice, and old pan-ERR agonist treated mice. We identified a subset of genes and proteins with V2 highest loadings (threshold was arbitrarily set to 0.04 or lower than −0.04) for further analysis.

### Western blotting

Western blotting was performed as previously described ^25, 36–38^. Equal amount of total protein was separated by SDS-PAGE gels and transferred onto PVDF membranes. After HRP-conjugated secondary antibodies, the immune complexes were detected by chemiluminescence captured on Azure C300 digital imager (Dublin, CA) and the densitometry was performed with ImageJ software. Primary antibodies used for western blotting were listed in **Supplementary Table 1**.

### Cytokine arrays

Cytokines in 200 microgram kidney lysate (pooled from 5-6 samples with equal amount of protein) were detected with the Proteome Profiler Array (ARY028; R&D Systems) according to manufacturer’s instructions.

### Statistical analysis

Results are presented as the means ± SEM for at least three independent experiments. Following the Grubbs’ outlier test, the data were analyzed by ANOVA and Newman–Keuls tests for multiple comparisons or by *t* test for unpaired data between two groups (Prism 6, GraphPad, San Diego, CA). Statistical significance was accepted at the *P*<0.05 level.

### Study approval

Animal studies and relative protocols were approved by the Animal Care and Use Committee at the Georgetown University. All animal experimentation was conducted in accordance with the Guide for Care and Use of Laboratory Animals (National Institutes of Health, Bethesda, MD).

## RESULTS

### ERRα and ERRγ expression is decreased in the aging human kidney

In a previous study, we found decreased expression of ERRα and ERRγ in the aging mouse kidney and expression was reversed by the dual FXR-TGR5 agonist INT-767 or CR. Further, increased ERR expression correlated with increased mitochondrial biogenesis and function in the treated aging kidneys ^12^. In light of the role of ERR in mitochondrial biogenesis, we determined whether decreased expression also occurs in the aging human kidney. We performed immunohistochemistry with human kidney sections from young and old individuals. The results indicate that both ERRα and ERRγ are expressed in renal tubules and that their expression levels were markedly decreased in aging human kidney (**Figure 1A**).

**Figure 1.**
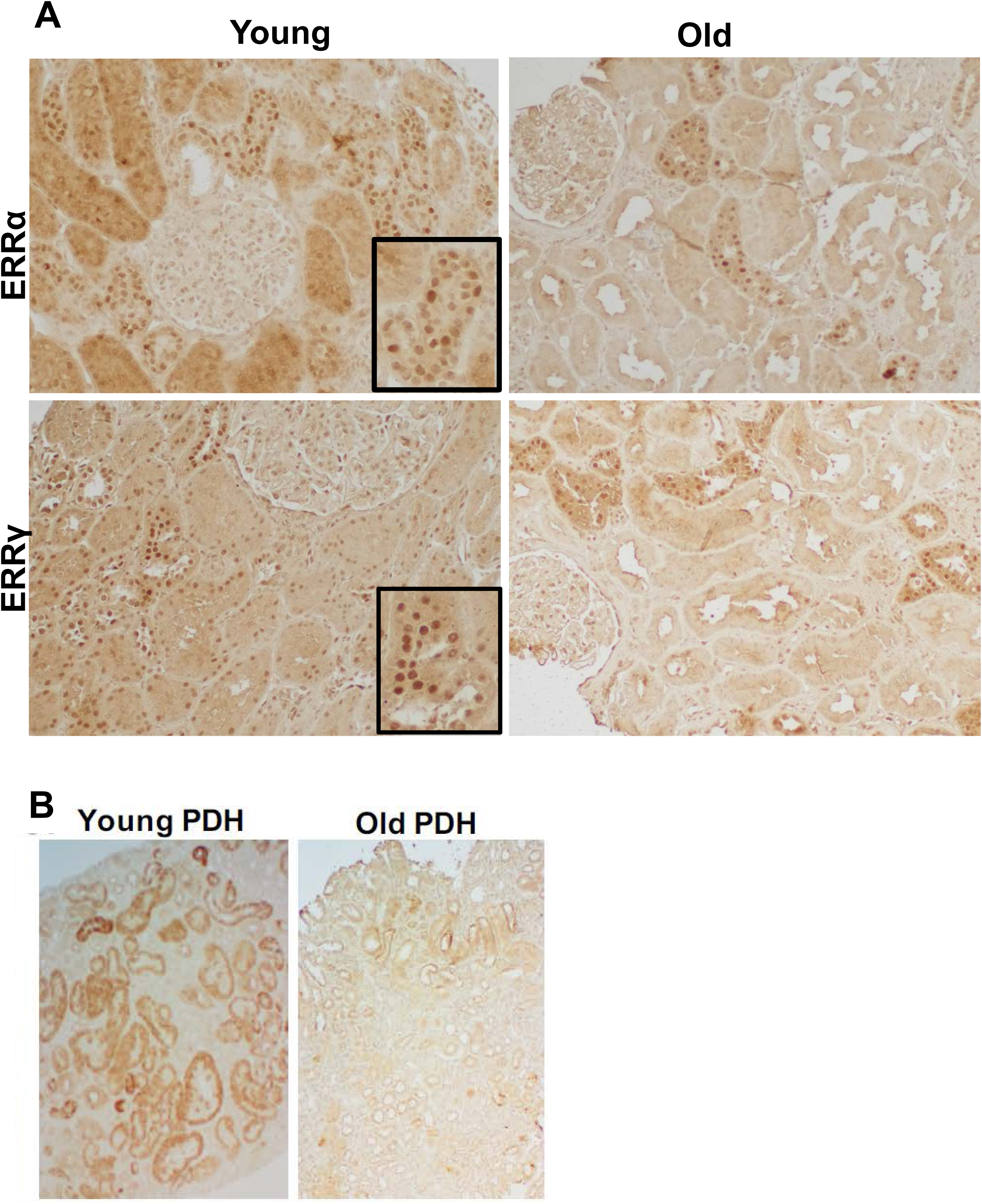
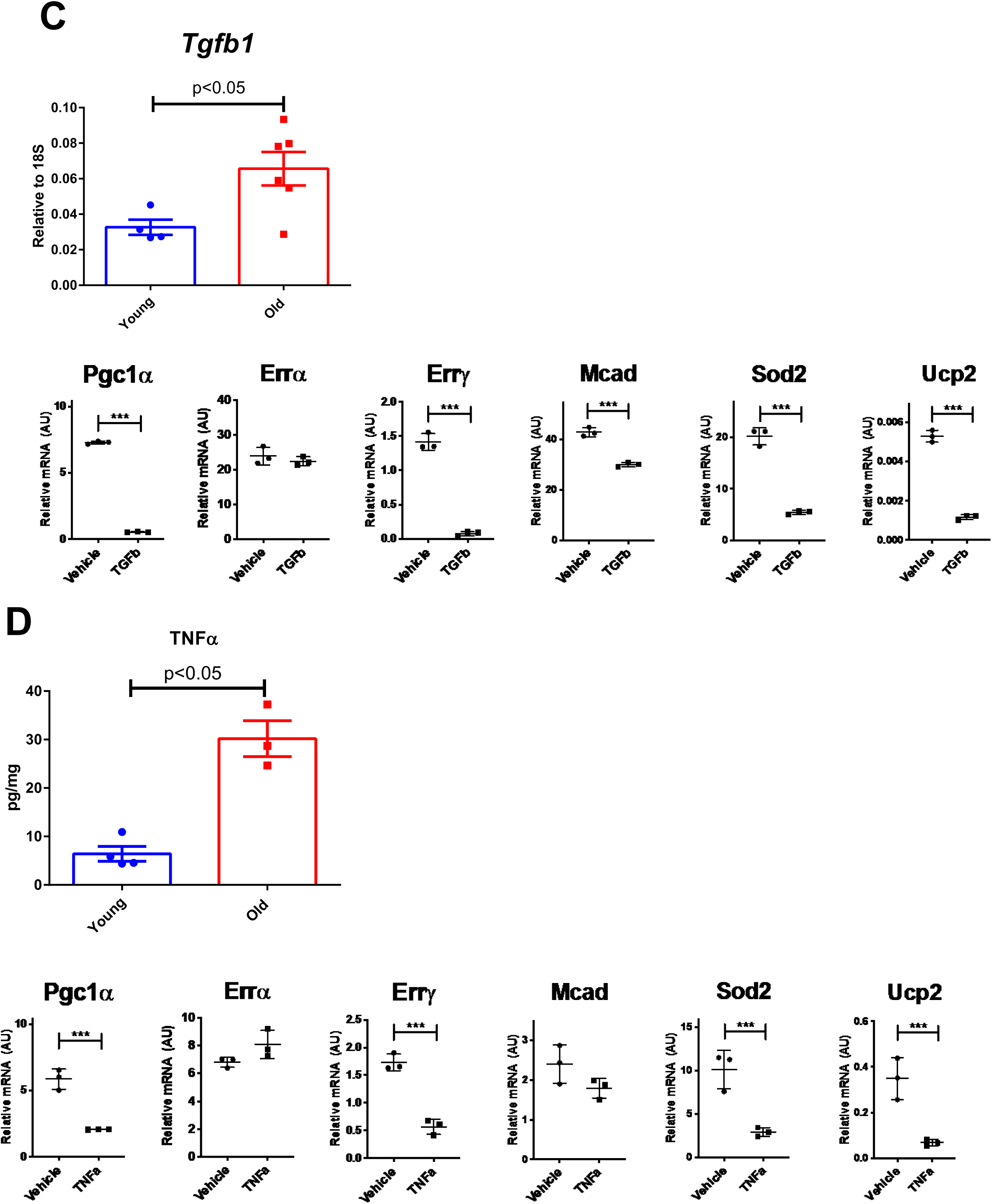
ERRα, ERRγ, and PDH expression is decreased in the aging human kidney. **A)** ERRα and ERRγ immunohistochemistry on kidney sections from young and old subjects. The nuclear positive staining for ERRα or ERRγ is decreased in old kidney sections. Box shows magnified image. **B)** PDH immunohistochemistry on kidney sections from young and old subjects. PDH staining is decreased in old kidney sections. **C-D)** TGF-β mRNA **C)** and TNFα protein expression **D)** were increased in the aging kidney. n=4-6 samples per group. TGF-β 1 **C)** and TNF -α **D)** treatment of human primary proximal tubule cells (PTEC) significantly decreased expression levels of ERRγ mRNA but not ERRα. PGC1α, MCAD, SOD2, and UCP2 gene expression were also significantly decreased in TGF-β treated cells **C)**. TNF also had a similar effect to decrease PGC1α, ERRα, SOD2, and UCP2 expression **D)**. N=3 for each group. *** p<0.001.

Since ERRs are important modulators of mitochondrial biogenesis, we also stained the human kidney sections for the mitochondrial pyruvate dehydrogenase (PDH) e2/e3 and found a marked decrease in PDH immunostaining in the aging human kidney samples (**Figure 1B**).

### TGF-**β** and TNF-**α** mediate decreases in ERR**γ** in cultured human proximal tubular epithelial cells

Transforming growth factor-β (TGF-β) and tumor necrosis factor-α (TNF-α) expression are increased in the aging kidney (**Figure 1C-D**). Thus, we determined whether TGF-β and TNF-α at least in part mediate the decreased expression of ERR in the kidney. We found that in cultured human primary proximal tubular cells (PTEC), TGF-β significantly decreased expression level of ERRγ mRNA while ERRα was unaffected. Expression of PGC1α, MCAD (encoding medium chain acyl-CoA dehydrogenase), SOD2, and UCP2 were also decreased in TGF-β treated cells (**Figure 1C**).

TNF-α also had a similar effect to decrease PGC1α, ERRγ, SOD2, and UCP2 expression (**Figure 1D**).

### ERRα and ERRγ RNA distribution in the mouse kidney

To determine where ERRα and ERRγ mRNA are expressed in the mouse kidney, we performed single nuclei RNAseq ^29, 30^. With 3,000-5,000 nuclei sequenced at 100k read depth, 12 expression clusters were identified and assigned to major cell types in the mouse kidney (**Figure 2A**). We found that ERRα was expressed in most of proximal tubules, intercalated cells and podocytes. ERRγ was detected in much less cells and mainly appeared in proximal tubules and intercalated cells (**Figure 2B**). Compared to young kidneys, S1/S2 segments of aging proximal tubules showed a decline in both ERRα and ERRγ expression (**Figure 2C**).

**Figure 2.**
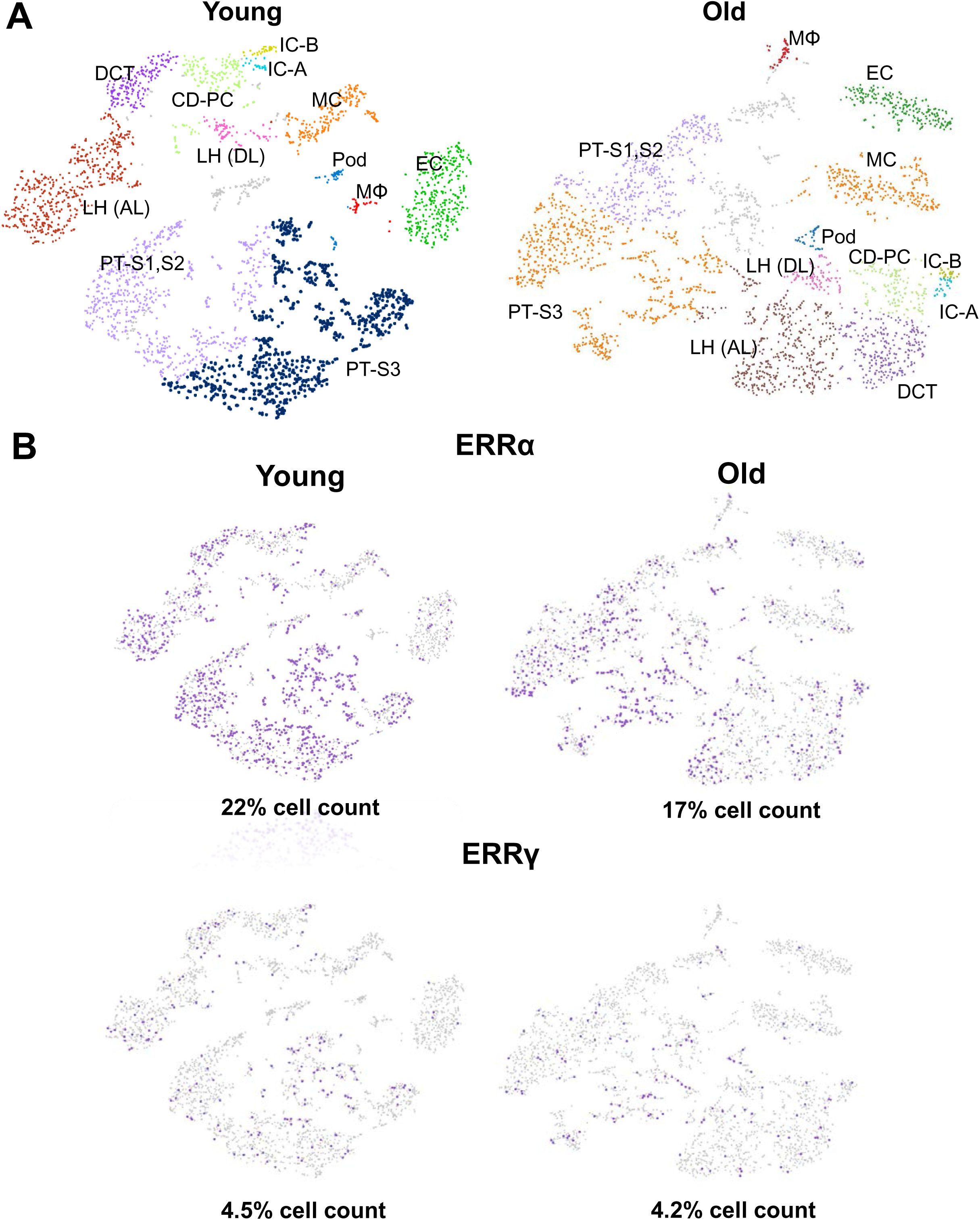
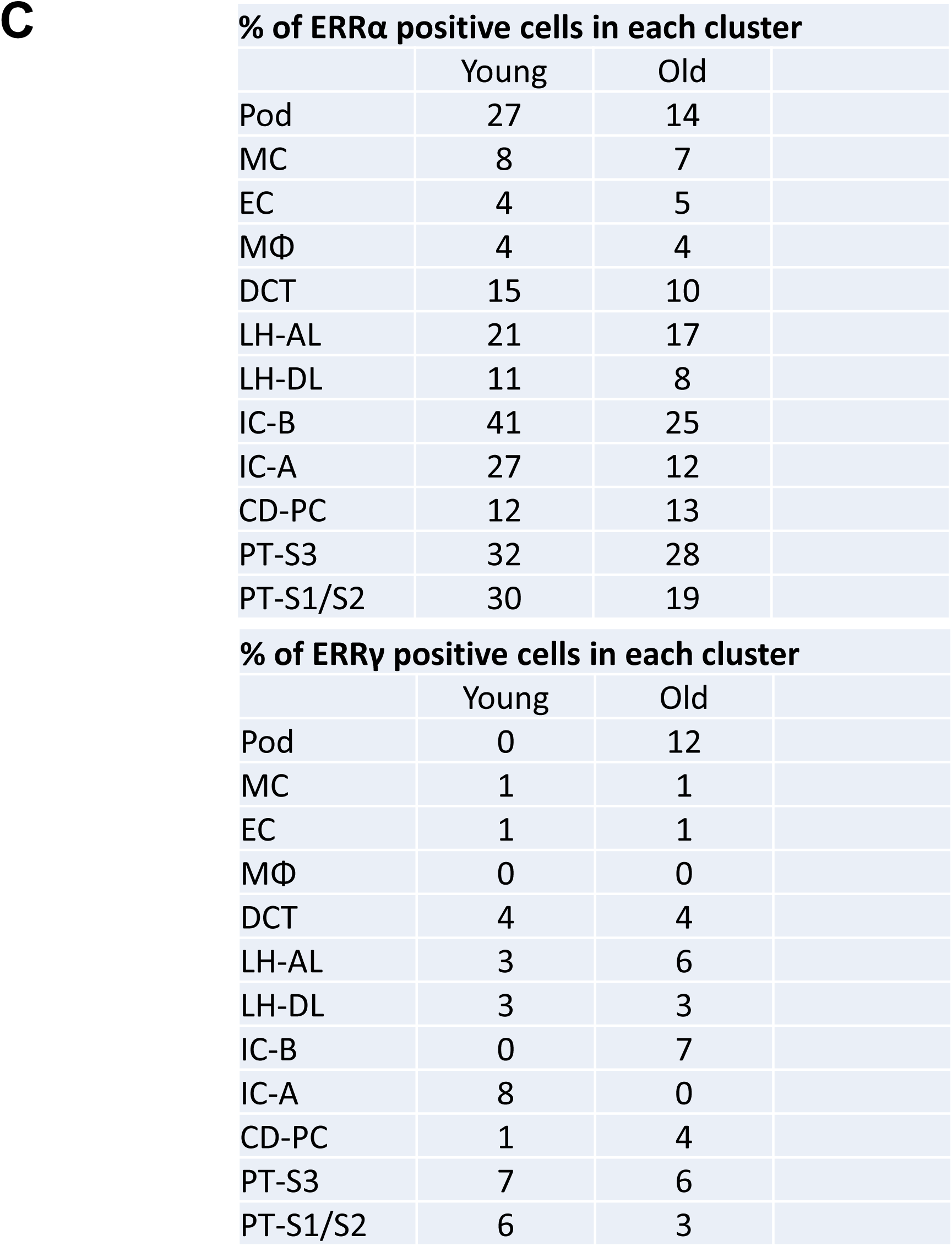
Single nuclei RNAseq of young and old kidneys. **A)** The t-distributed stochastic neighbor embedding (tSNE) shows that, with 100k read depth and 3000-5000 nuclei sequenced, we were able to identify 12 clusters and assigned them to major cell types known in the mouse kidney. **B**) We found most ERRα is expressed in proximal tubules, intercalated cells and podocytes. For ERRγ, proximal tubules and intercalated cells expressed most strongly. The cells with positive expression of ERRα or ERRγ were labeled purple. **C)** Table to show percentage of positive expressed cells in each cluster. Compared to the young kidneys, aging proximal tubules at S1/S2 show decline in ERR α and ERR γ expression. Pod: podocyte; MC: mesangial cell; EC: endothelial cell; Mφ : macrophage; DCT, distal convoluted tubule; LH, loop of Henle; IC: intercalated cell; CD-PC, collecting duct-principal cell; PT, proximal tubule.

### Pan-ERR agonist treatment improves the age-related kidney injury

We treated the aging mice with a recently available pan-ERR agonist (SLUPP-332). We found that 2-month treatment of 21-month-old mice significantly improved the age-related albuminuria, and decreased kidney weight (**Figure 3A**). The decrease in albuminuria is likely related to improved podocyte function as protein expression of the podocyte marker NPHS2 (podocin) is increased after the treatment (**Figure 3B**). In addition, mRNA expression in profibrotic markers (TGF-β PAI-1 and Col IV), monocyte/macrophage marker (F4/80) and tubular injury marker neutrophil gelatinase-associated lipocalin (NGAL) were decreased with the pan-agonist treatment (**Figure 3C**). Finally, with cytokine profiling we found the kidney injury-related proteins, Ngal, kidney injury marker-1 (Kim1), osteopontin and CC-motif ligand chemokine-21 (Ccl21) were increased in the aging kidney and partially normalized by pan-ERR agonist treatment (**Figure 3D**).

**Figure 3.**
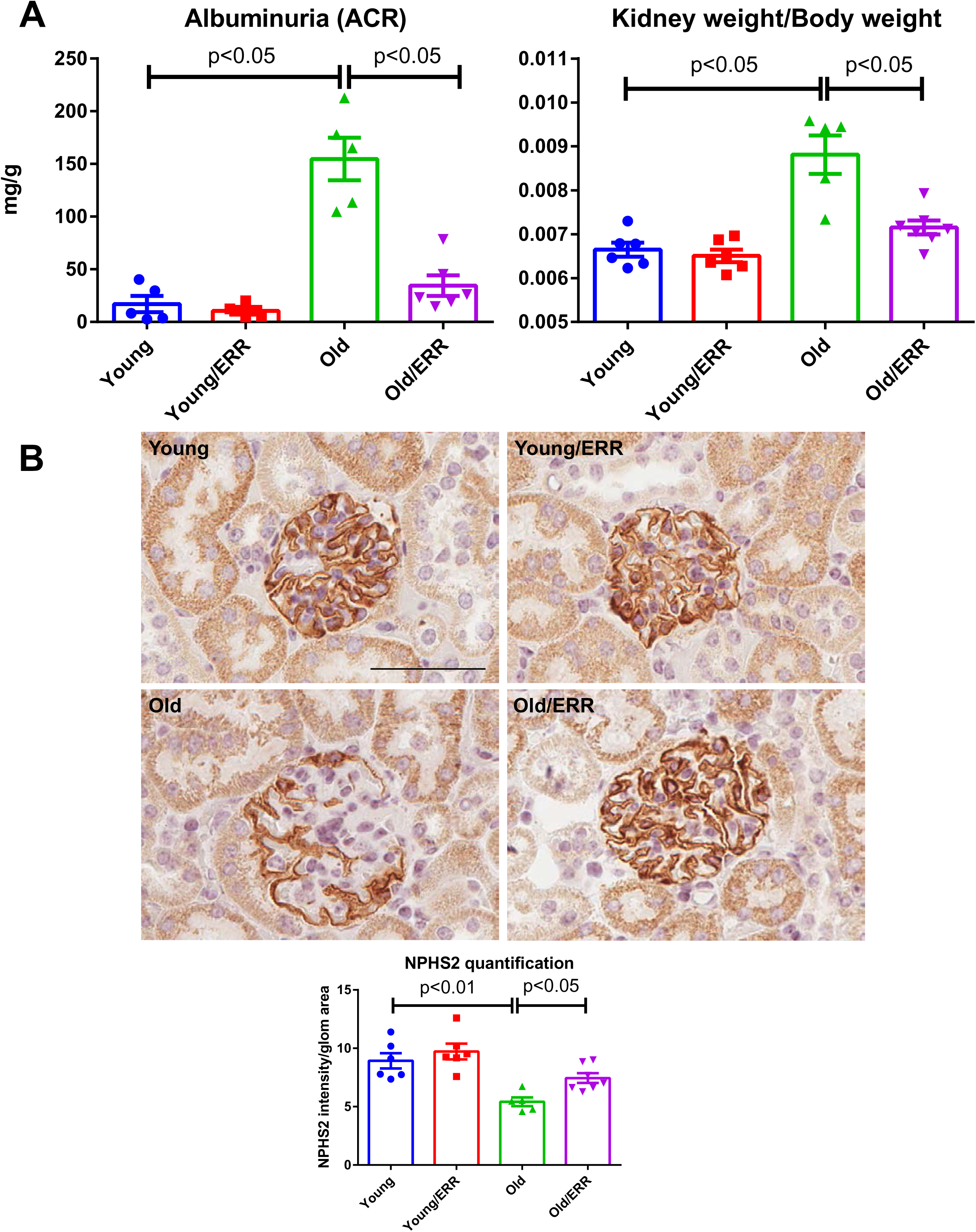
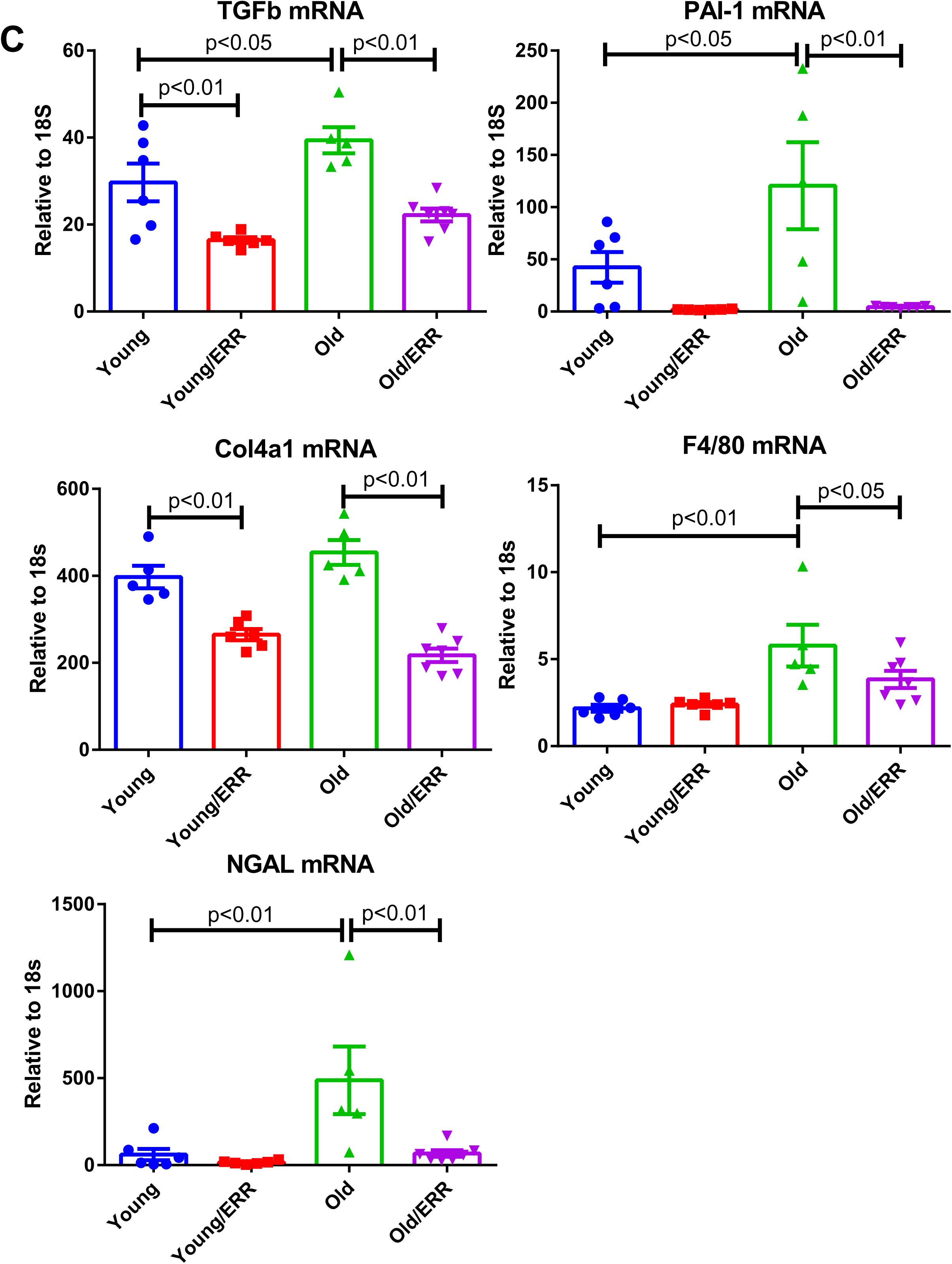
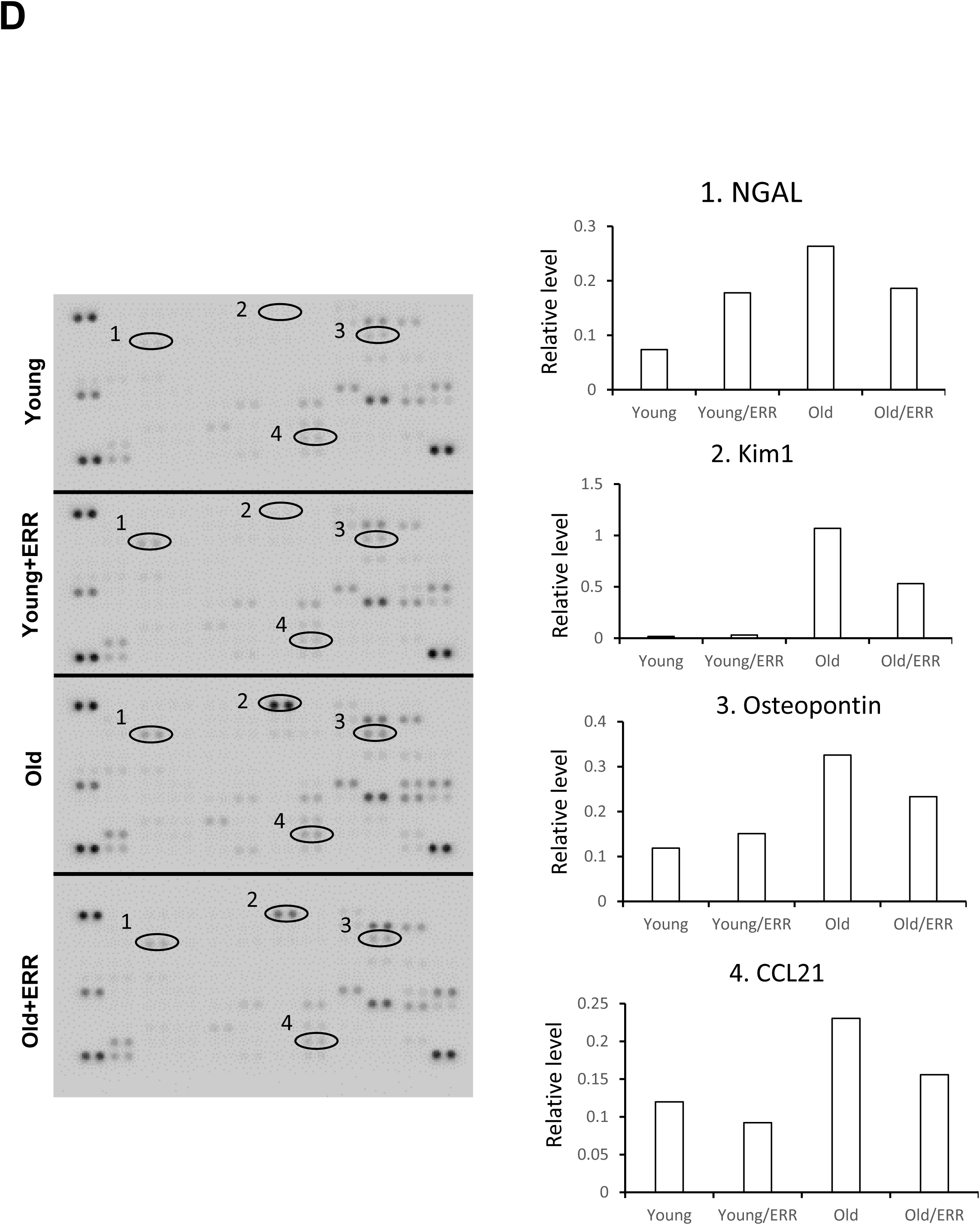
Pan-ERR agonist improved age-related renal injury. **A)** Albuminuria and kidney weight (normalized by body weight) were increased in aging kidneys and normalized with treatment. N=5-6 for each group**. B)** NPHS2 (podocin) immunohistochemistry on kidney sections, labeling podocytes. Pan-ERR agonist treatment reversed the decreased NPHS2 staining in aging kidneys. Scalar bar: 50 µm. N=5-6 for each group**. C)** Kidney injury markers TGF-β, PAI-1, Col4a1, F4/80, and NGAL mRNA expression were increased in aging kidneys but decreased with ERR treatment. N=5-6 for each group. **D)** Cytokine array showed 4 major spots that correspond to kidney injury markers Ngal, Kim1, Osteopontin and Ccl21, with increased expression in the aging kidney and decreased expression following the pan agonist treatment. Each group was one sample pooled from n=4 animals.

### Pan-ERR agonist treatment modulates mitochondrial metabolism and inflammation in the aging kidney

To assess the actions of the pan-ERR agonist, we examined ERRα, ERRβ and ERRγ mRNA abundance. Although they were all decreased in the kidneys of aging mice, treatment with the ERR pan-agonist induced significant increases in their expression to levels observed in the young mice (**Figure 4A**).

**Figure 4.**
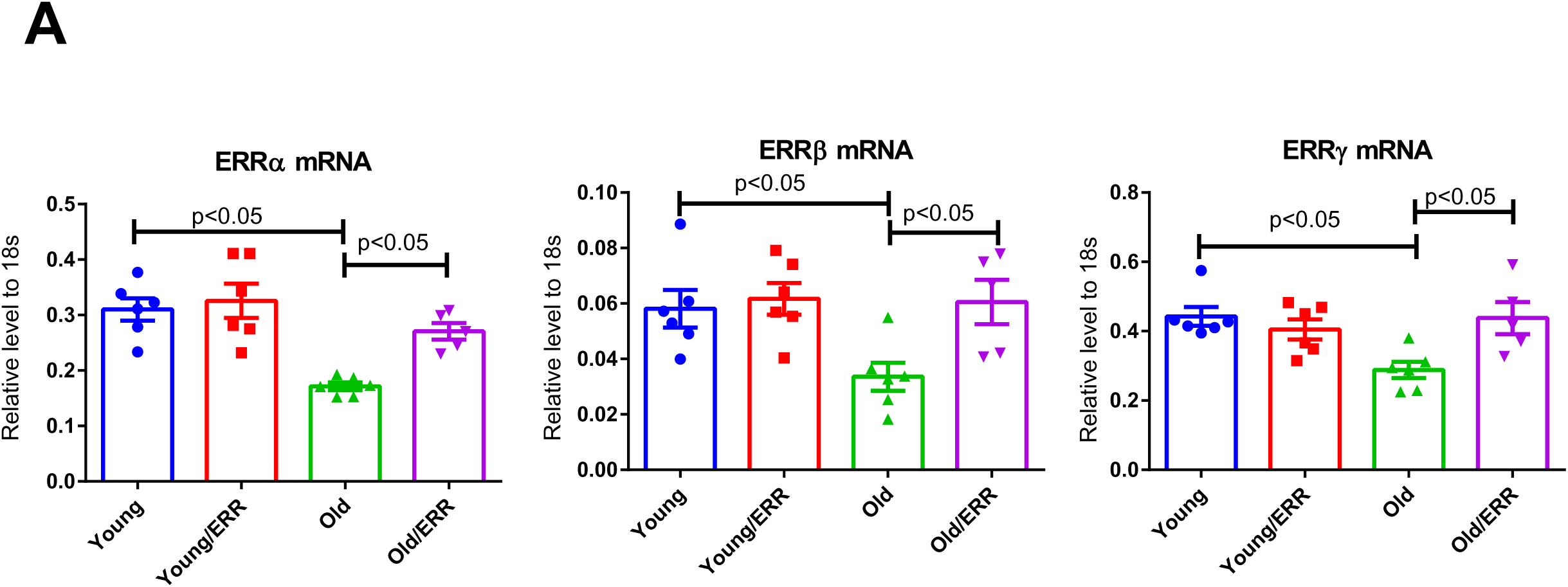
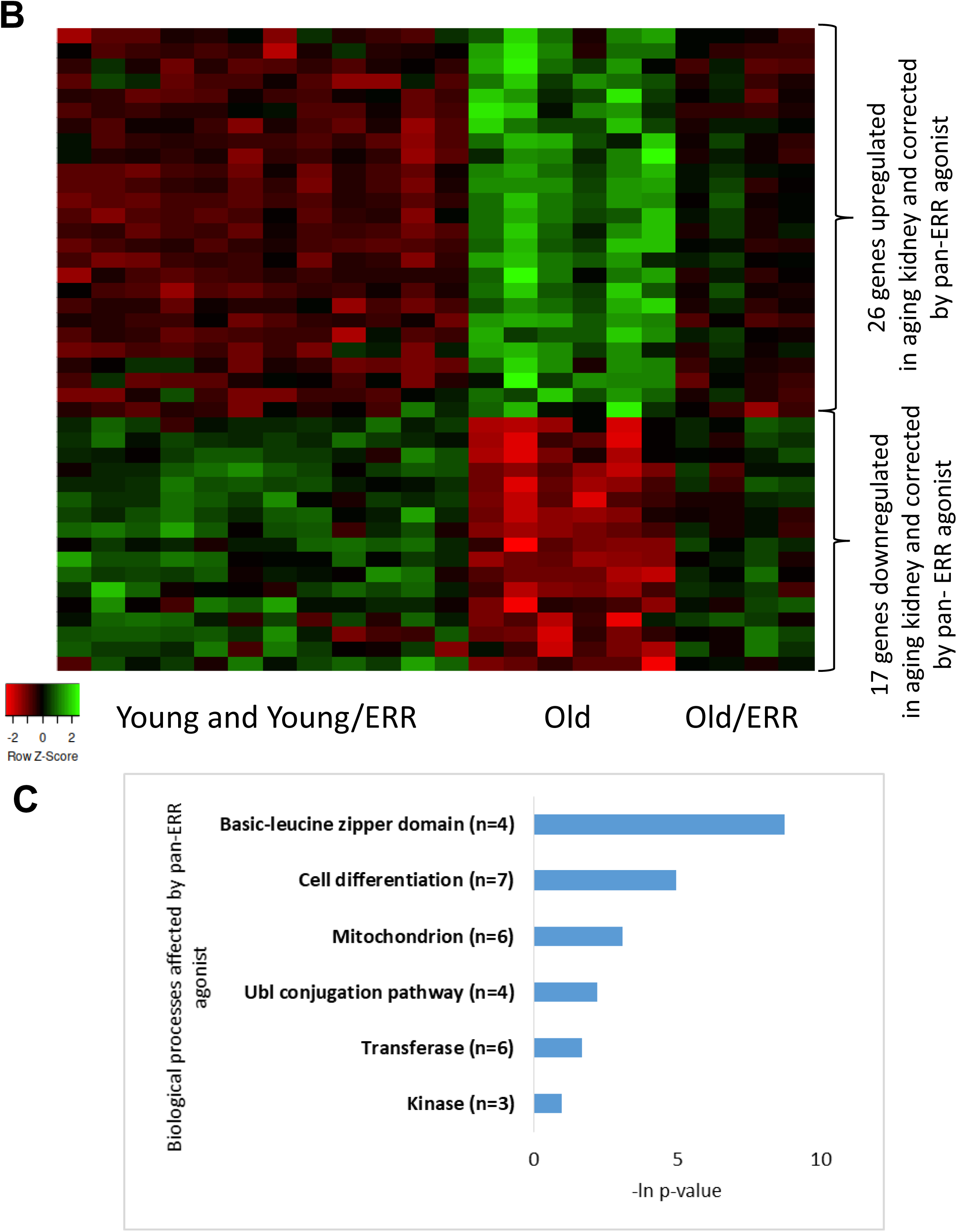
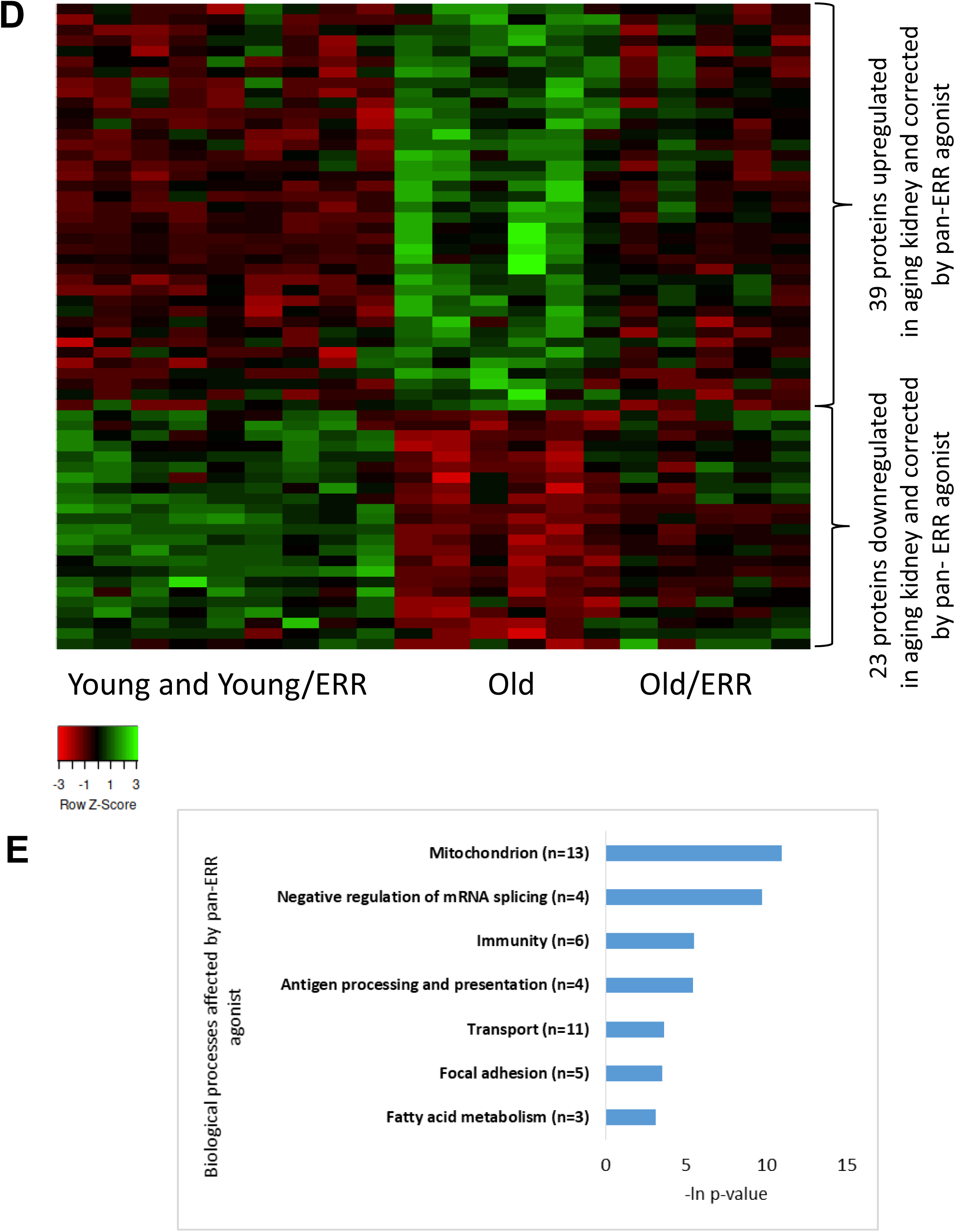
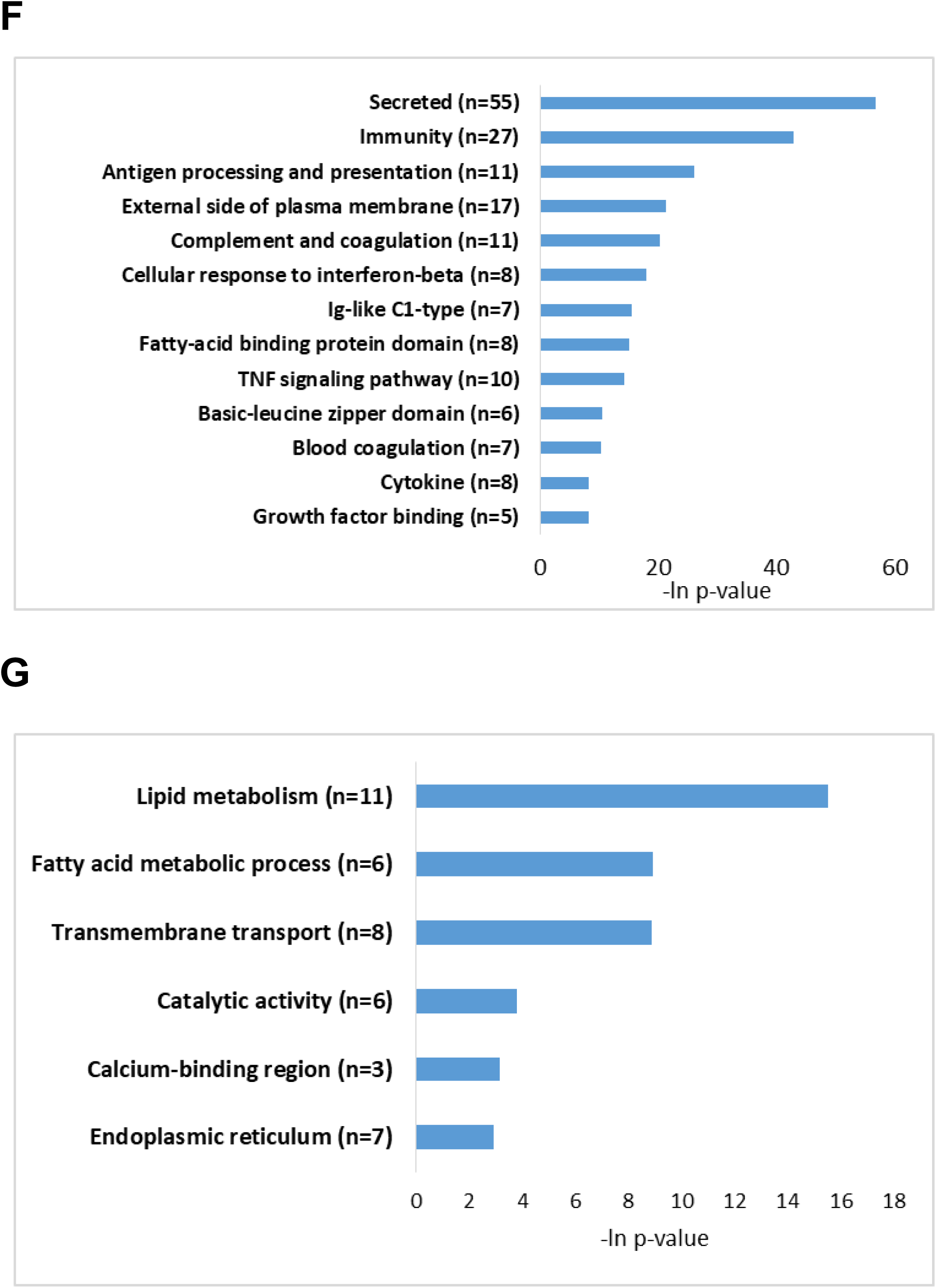
RNAseq and proteomics of kidney from old mice treated with vehicle or the pan-ERR agonist. **A)** ERRα, ERRβ and ERRγ mRNA expression were decreased in aging mouse kidneys. Treatment with the pan ERR agonist restored the mRNA levels of ERRα, ERRβ and ERRγ to levels seen in young kidneys. N=5-6 for each group. **B)** Heat map showing expression patterns of genes differentially expressed in kidneys of old mice treated with vehicle compared to kidneys of old mice treated with pan-ERR agonist. The heat map indicates up-regulation (green), down-regulation (red), and unaltered gene expression (black). The columns represent individual samples. **C)** Functional pathway enrichment analysis of differentially expressed proteins in kidneys of old mice treated with vehicle compared to kidneys of old mice treated with pan ERR agonist. The y-axis shows significantly enriched pathways. The x-axis indicates p-value of enrichment of the given pathway. **D)** Heat map showing expression patterns of proteins differentially expressed in kidneys of old mice treated with vehicle compared to kidneys of old mice treated with pan ERR agonist. The heat map indicates up-regulation (green), down-regulation (red), and unaltered gene expression (black). The columns represent individual samples. **E)** Functional pathway enrichment analysis of differentially expressed proteins in old mice treated with vehicle compared to kidneys of old mice treated with pan ERR agonist. The y-axis shows significantly enriched pathways. The x-axis indicates p-value of enrichment of the given pathway. **F)** Functional pathway enrichment analysis of subset of genes and proteins identified with O2PLS analysis as upregulated in kidneys of old mice compared to kidneys of young mice and that were downregulated by ERR treatment in old mice. The y-axis shows significantly enriched pathways. The x-axis indicates p-value of enrichment of the given pathway. **G)** Functional pathway enrichment analysis of subset of genes and proteins identified with O2PLS analysis as downregulated in kidneys of old mice compared to kidneys of young mice and that were upregulated by ERR treatment in old mice. The y-axis shows significantly enriched pathways. The x-axis indicates p-value of enrichment of the given pathway.

Next, we performed bulk mRNA sequencing analysis to determine which pathways are affected by aging and are restored by pan-ERR agonist. We first investigated the molecular changes in aging kidney. We found 448 upregulated and 463 downregulated genes in kidneys of aging mice compared to young controls (**Supplementary Figure 2A, Supplementary Table 2**). Pathway enrichment analysis of upregulated genes revealed that inflammation-related pathways were highly significant. The inflammation-related pathways included ‘Inflammation mediated by chemokine and cytokine signaling pathway’, ‘EGF receptor signaling pathway’, ‘Toll receptor signaling pathway’, and ‘TGF-beta signaling pathway’ which were all previously reported as upregulated in aging kidney^39, 40^ (**Supplementary Figure 2B**).

In addition to the inflammation, apoptosis and cell-cycle regulating pathways were also enriched with upregulated genes in the aging kidney (**Supplementary Figure 2B**). The cell-cycle dysregulation is a known phenomenon in the aging and diseased kidney and might indicate cell senescence ^41^. Indeed, when we analyzed the RNA-seq data in a supervised manner identifying senescence associated genes, we found that in aging kidney 29 senescence associated genes were upregulated and 10 were downregulated (**Supplementary Figure 2C, Supplementary Table 3**). In particular, we found that p53 (Trp53), p65 (Rela), and p21(Cdkn1a) which are the major indicators of senescent cells were highly expressed in aging kidney. The downregulated genes in aging kidney were enriched in mitochondrial and metabolic processes (**Supplementary Figure 2D**). Dysregulated mitochondrial functioning has been widely studied in aging kidneys^42^. Recently, several studies have shown that damaged mitochondria may trigger inflammation via cGAS-STING pathway in kidney disease ^43, 44^. TFAM is a key regulator of mitochondrial gene expression ^45^ and is crucial for maintaining mtDNA structure, transcription, and replication^46^. Deletion of Tfam leads to mtDNA escape into the cytoplasm and activation of the innate immune pathway through cGAS-STING activation^44^. Our mRNA sequencing data indicates that Tfam mRNA levels are decreased in aging kidney (**Supplementary Figure 2E**). Overall, we found that in aging kidney there is a dysregulation of mitochondrial and immune processes.

We next examined the effect of pan-ERR agonist treatment on gene expression. Pan-ERR agonist treatment mitigated several of the abovementioned pathways (**Figure 4B-C, Supplementary Table 4**).

In addition to mRNA sequencing, we performed mass-spec proteomics. Because the concordance of proteomics and RNA transcriptomics is known to be weak ^47, 48^, we expected the proteomics analysis to reveal additional processes disrupted in aging and attenuated by pan-ERR agonist treatment. We found that, on the protein level, mitochondria-related, oxidation reduction processes, peroxisome, and metabolic pathways were dysregulated in aging kidney and correlated with pathways found on mRNA level. Furthermore, several additional dysregulated pathways were identified from proteomics analysis in aging kidney. These additional processes included calcium binding region, blood coagulation, and biotin/lipoyl attachment processes (**Supplementary Figure 2F-G, Supplementary Table 5**). The processes of mitochondrion, immune-related pathways, transport regulation, focal adhesion, and fatty acid oxidation were improved by pan-ERR agonist treatment (**Figure 4D-E, Supplementary Table 6**).

It has been suggested that combining several layers of omics data yields clearer understanding of complex biological phenomena than evaluation of each layer separately ^49, 50^. Thus, applying the two-way orthogonal Partial Least Squares (o2PLS) ^51^, we found that one of the components (V2) reflects the expected relation between the biological groups under investigation. Namely, V2 component profile is associated with pan-ERR agonist effect of alleviating the dysregulation of genes and proteins in aging kidneys (**Supplementary Figure 3**). Genes and proteins whose expression correlates with V2 component profile were identified as an interrelated molecular network associated with overall changes in aging and pan-ERR agonist treatment (see Methods, **Supplementary Table 7**). We found that the subset of genes and proteins upregulated in aging kidney and downregulated by pan-ERR agonist treatment were enriched with immune related processes such as antigen processing and presentation, complement and coagulation cascades, Ig-like C1-type domain protein activation, and TNF signaling pathway (**Figure 4F**). The subset of genes and proteins downregulated in aging kidney and enhanced by pan-ERR agonist treatment was enriched by processes associated with metabolism such as lipid metabolic processes, fatty acid metabolism, and catalytic activity (**Figure 4G**).

Taken together, our analyses of transcriptomic and proteomic changes as well as integrated analyses of transcriptome-proteome associations point to downregulation of mitochondrial metabolism and upregulation of inflammatory processes in aging kidney. Theses analyses also showed that pan-ERR agonist treatment was effective in attenuating these age-related dysregulations. Thus, our analyses correlate with previous studies to show that the hallmarks of aging kidneys are decreased mitochondrial function and increased inflammation ^52, 53^. We next determined whether treatment with the pan-ERR agonist improved these specific defects in the aging kidneys.

### Pan-ERR agonist treatment restored mitochondrial function in aging kidneys

The canonical function of ERR is to induce mitochondrial biogenesis. We found that the pan-ERR agonist increased expression of the master mitochondrial biogenesis regulators PGC1α and PGC1β in aging kidneys. Consistently, mitochondrial transcription factor *Tfam1* expression was decreased in the aging kidney and increased with the pan-ERR agonist. As a result, the mitochondrial DNA/nuclear DNA ratio was increased in aging kidneys following treatment, indicative of increased mitochondrial biogenesis (**Figure 5A**). The increased mitochondrial biogenesis was accompanied by increased expression of genes related to the mitochondrial ETC complexes, such as complex I subunit *Ndufb8*, complex II subunit *Sdhc*, complex III subunit *Uqcrb*, complex IV subunit *Cox6a2*, and complex V subunit *Atp5b* (**Figure 5B**). Native blue gel analysis further showed increased quantities of assembled mitochondrial complexes after the treatment (**Supplementary Figure 4**). This is consistent with the increased gene expression of enzymes in TCA cycle, such as *Pdhb*, *Mdh1*, *Idh3b*, and *Sucla2* (**Figure 5C**). Since the TCA cycle can produce NADH which is required for complex I driven mitochondrial oxidative phosphorylation, we further determined fractional intensity of free NADH with phasor approach to fluorescence lifetime imaging microscopy (FLIM) ^54, 55^. Analysis of the FLIM images showed that there was decreased free NADH fraction in aging kidneys, which suggests a lower capacity of NADH regeneration in aging kidney, in agreement with the downregulation of TCA cycle enzyme expression ^56^. Treatment with the pan-ERR agonist shifted the free NADH fraction in the aging kidney towards that seen in the young kidneys as shown in cumulative plots for all samples ^55, 57^ (**Figure 5D**).

**Figure 5.**
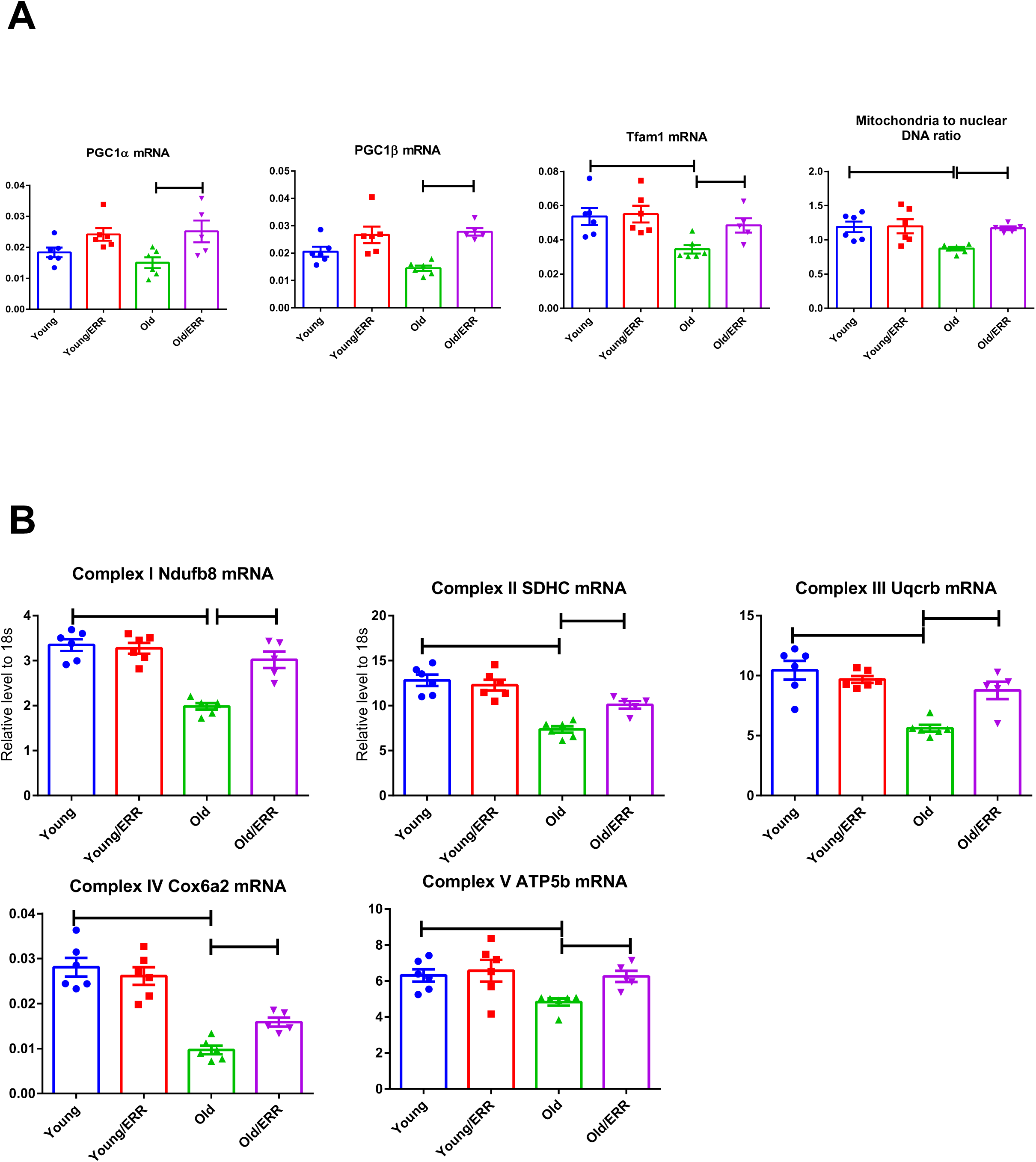
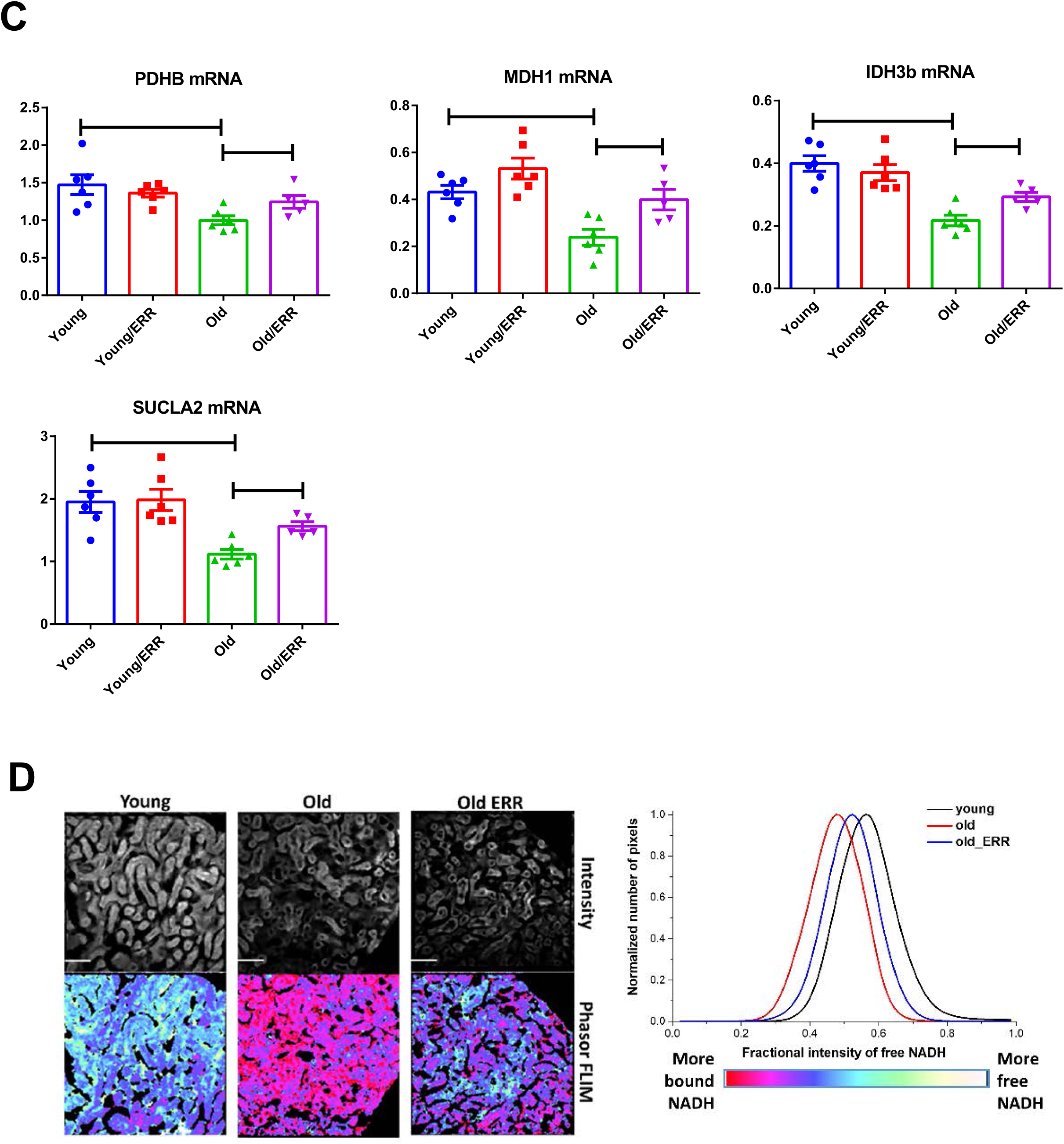
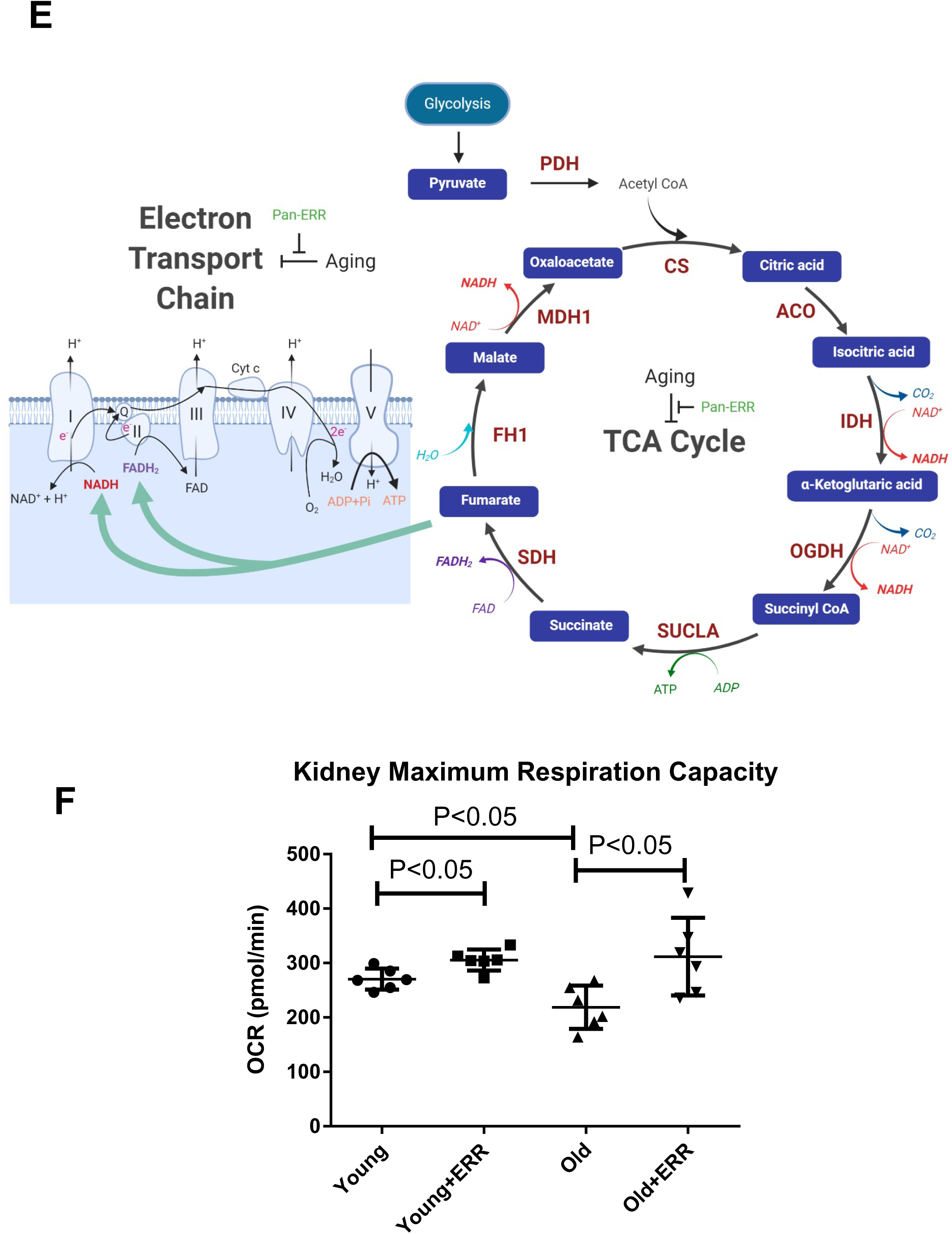
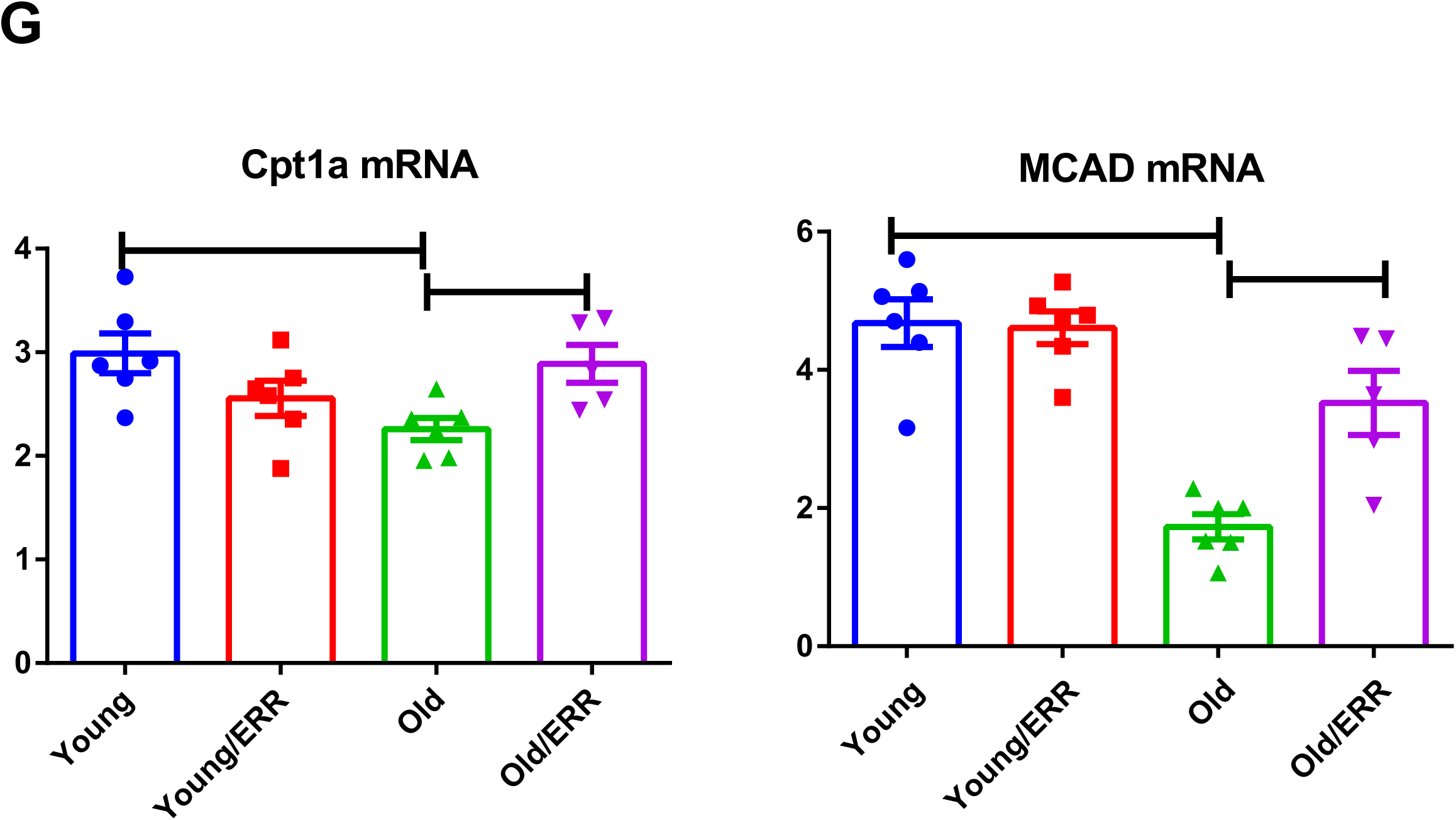
Pan-ERR agonist treatment restored mitochondrial function in aging kidneys. **A)** Pan-ERR agonist treatment in old mice increased the expression of PCG1α and PGC1β, coregulators of ERRs and mediators of mitochondrial biogenesis. The expression of the mitochondrial transcription factor Tfam1 was decreased in the kidneys of old mice and pan ERR agonist treatment restored it to levels seen in young mice. Mitochondria to nuclear DNA ratio was decreased in the kidneys of old mice and pan ERR agonist treatment restored it to levels seen in young mice, indicative of increased mitochondrial biogenesis. N=5-6 for each group. **B)** mRNA expression levels of subunits for mitochondrial electron chain complex (ETC) was decreased in the kidneys of old mice and pan ERR agonist treatment restored it to levels seen in young mice, indicative of improvement in ETC. N=5-6 for each group. **C)** mRNA expression levels of the TCA cycle enzymes were decreased in the kidneys of old mice and recovered following pan-ERR agonist treatment to levels seen in young mice, indicative of restoration of the TCA cycle. One of the TCA cycle intermediates, succinic acid showed increased level in the kidneys of old mice treated with the pan ERR agonist. N=5-6 for each group. **D)** Autofluorescence intensity (NADH channel) (top) and Phasor mapped FLIM image (bottom) on the kidney section. Scale bar = 100 µm. The phasor mapped FLIM images were pseudo colored based on the fractional intensity of the free NADH as shown in the color scale. The fractional intensity contributions were calculated by resolving the phasor signatures from these images in between the phasor positions of free and bound NADH. Fractional intensity of NADH plot showed that there was decreased free NADH fraction in aging kidneys and pan-ERR agonist treatment increased the free NADH fraction. N=5-6 for each group. **E)** Interrelationship of the TCA cycle and ETC. **F)** Maximum respiration capacity in mitochondria isolated from the kidneys showed a significant impairment in old mice, which is restored to levels seen in young mice after treatment with the pan ERR agonist. N=5-6 for each group. **G)** The fatty acid β-oxidation enzymes *Cpt1a* and *Mcad* mRNA levels were decreased in the kidneys of old mice, which were restored to levels seen in young mice after treatment with the pan ERR agonist. N=5-6 for each group.

The interrelationship between the ETC and TCA cycle is illustrated in **Figure 5E**. The changes in ETC complexes and TCA cycle resulted in increased maximum respiration capacity in mitochondria isolated from treated aging kidneys (**Figure 5F**). In addition, mRNAs encoding enzymes that mediate mitochondrial fatty acid β *Cpt1a* and *Mcad*, were upregulated by the pan-ERR agonist (**Figure 5G**), suggesting that ERR agonism promotes mitochondrial fatty acid β

### Pan-ERR agonist treatment altered mitochondrial dynamics in aging kidneys

Transmission electron microscopy showed alterations in the mitochondria of aging kidneys, including decreases in the area, perimeter, and minimum Feret diameter in the aging kidneys; these parameters were restored to levels seen in young kidneys upon treatment with the pan-ERR agonist (**Figure 6A**). Since these mitochondrial changes are reminiscent of alterations in mitochondrial fusion and fission, we also measured the expression of proteins that regulate mitochondrial fusion and fission. Mitofusin 2 (Mfn2) is found in the outer membrane that surrounds mitochondria and participates in mitochondrial fusion ^58^. There was a significant decrease in Mfn2 in the aging kidney, and the ERR pan agonist increased the Mfn2 protein abundance in both the young and the old kidneys, fully restoring levels in the old kidneys to levels of the untreated young kidneys (**Figure 6B**).

**Figure 6.**
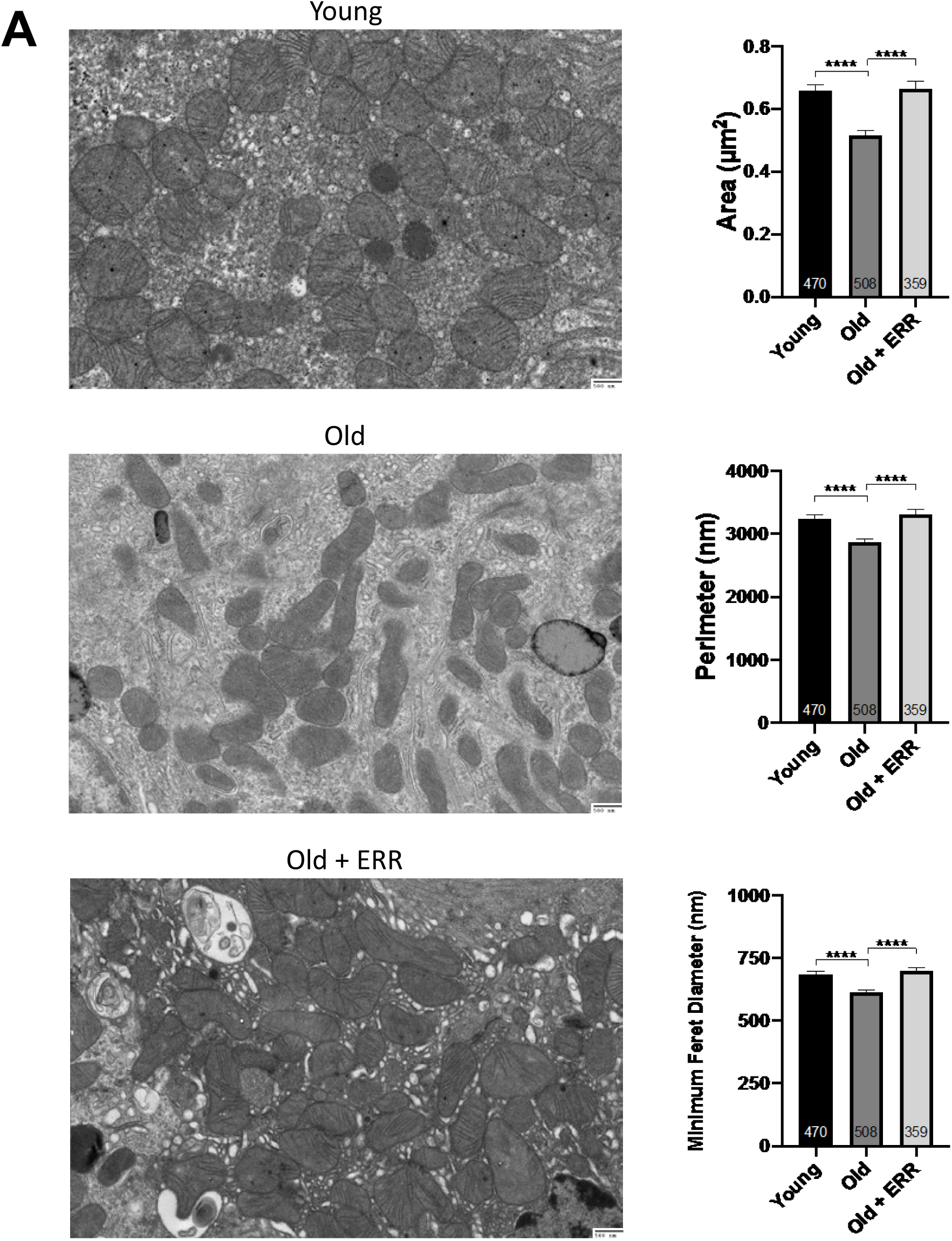
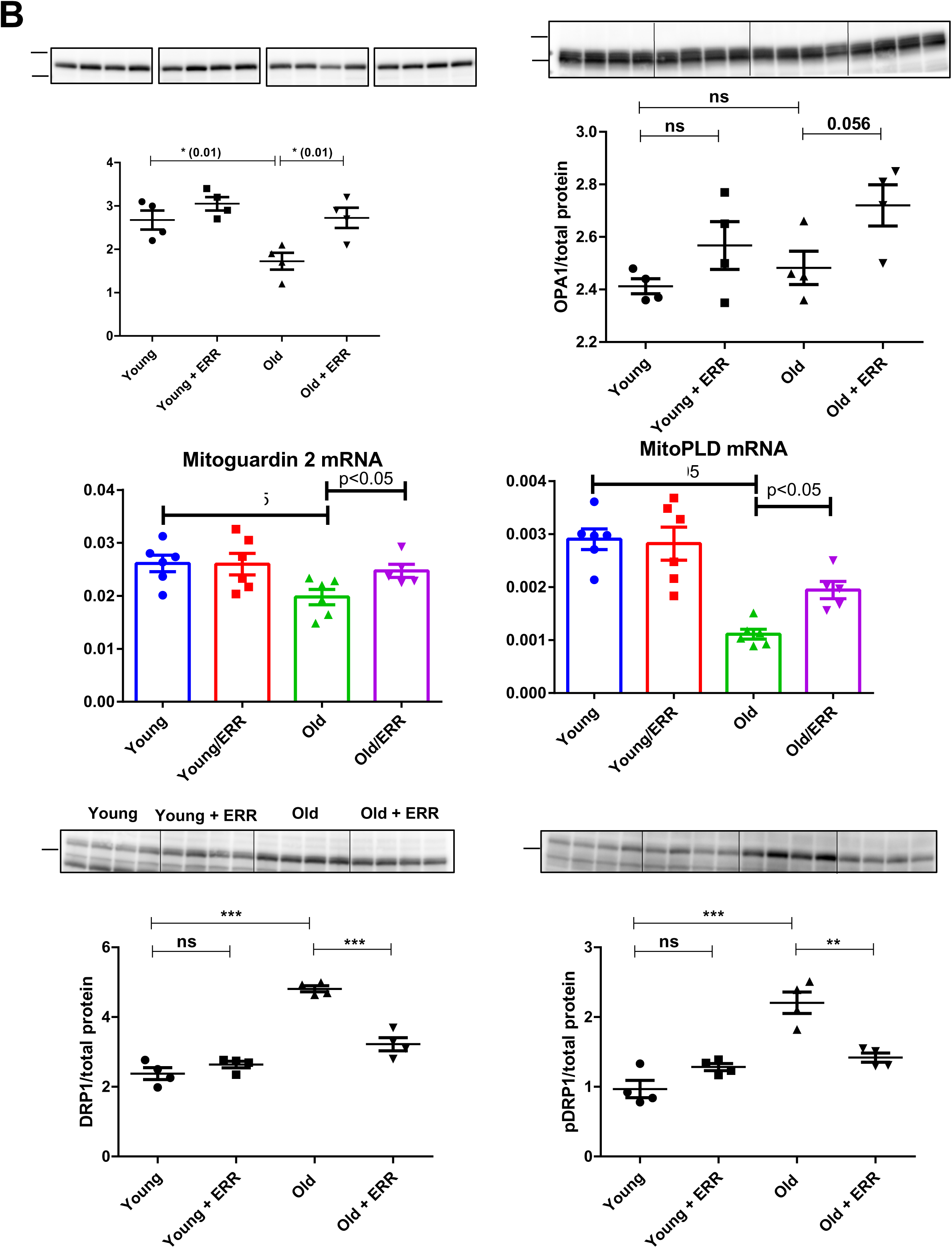
Pan-ERR agonist treatment altered mitochondrial dynamics in aging Kidneys. **A)** Transmission electron microscopy showed alterations in the mitochondria of aging kidneys including decreases in the area, perimeter, and minimum Feret diameter, which were restored to levels seen in young kidneys upon treatment with the pan ERR agonist. N = 3-4 for each group. **B)** There was a significant decrease in mitofusin-2 protein abundance in the old kidneys and the pan-ERR agonist increased the protein abundance in both the young and the old kidneys, with the resulting levels in the old kidneys being the same as in the young kidneys. In contrast, there was no significant change in the protein level of Opa1 in the old kidneys. However, upon treatment with the pan-ERR agonist, there was a tendency for the protein level to increase in the aging kidneys. There were also significant decreases in mitoguardin 2 and MitoPLD mRNA levels in the aging kidneys that were normalized upon treatment with the pan-ERR agonist. In addition, there were significant increases in Drp1 and phospho-Drp1 protein in the kidneys of old mice, which were restored back to levels seen in young mice following treatment with the pan ERR agonist. N=5-6 for each group in mRNA level analysis. N=4 for each group in protein analysis.

There were changes in expression of other nuclear-encoded, mitochondrial-expressed mRNAs and proteins that would be congruent with restoration of mitochondrial function. Opa1 protein localizes to the inner mitochondrial membrane and helps regulate mitochondrial stability, energy output, and mitochondrial fusion ^59^. While there was no significant change in the protein level in the aging kidney, upon treatment there was a tendency for the protein level to increase in the aging kidneys (**Figure 6B**). In addition, there were also significant decreases in mitoguardin 2 (*Miga2*) and mitoPLD (*Pld6*) mRNA levels in the aging kidneys that were increased upon treatment with the pan-ERR agonist (**Figure 6B**). Mitoguardin 2 regulates mitochondrial fusion through mitoPLD ^60^. Drp1 is a member of the dynamin superfamily of proteins and is a fundamental component of mitochondrial fission ^58^. There were significant increases in Drp1 and phospho-Drp1 protein in the kidneys of aging mice, which were restored to levels seen in young mice following treatment with the pan-ERR agonist (**Figure 6B**).

### Pan-ERR agonist treatment decreased inflammation in aging kidneys

Mitochondria are immunogenic organelles and mitochondrial dysfunction generates several immunogenic molecules, including mtDNA^61^ and mtRNA^62^. The cyclic GMP-AMP synthase (cGAS)-stimulator of interferon genes (STING) has been reported as one of the innate immune receptors to be activated by mtDNA leaking into cytosol ^44, 63^. In the aging kidneys, we found increased expression of STING and cGAS mRNAs and proteins, which was significantly reversed by treatment with the pan-ERR agonist (**Figure 7A**). We also found similar changes with RNA sensors RIG-I/MDA5/LGP2 and other nucleic acid sensors, such as TLRs ^64, 65^ (**Figure 7B**). We further examined the downstream response and we found out that expression of mRNA encoding components in NFκB signaling pathway (*Rel, RelB, Nfkb2*, and p65 protein) were increased in the aging kidney, and reduced by pan-ERR agonist treatment (**Figure 7C**).

**Figure 7.**
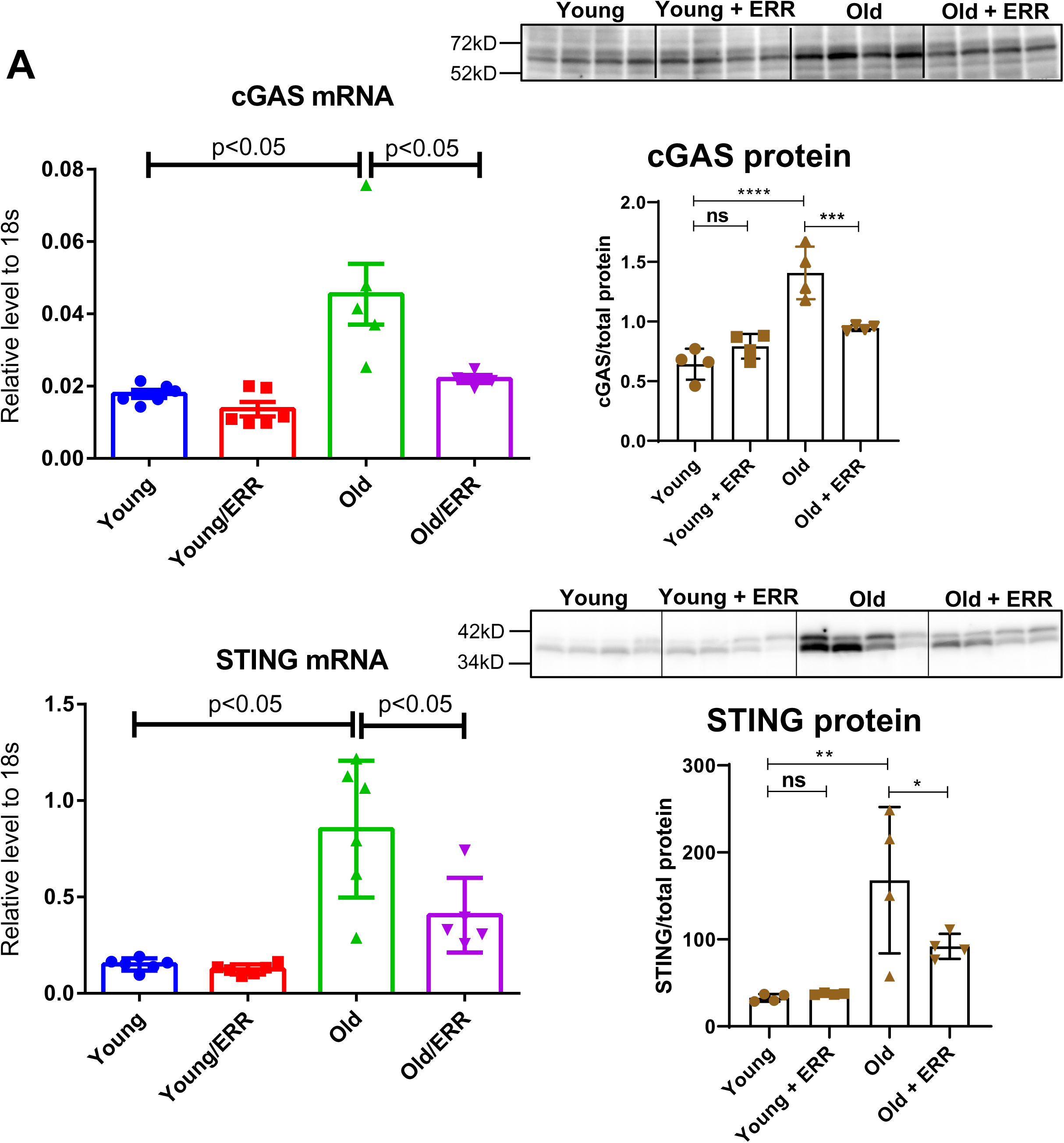
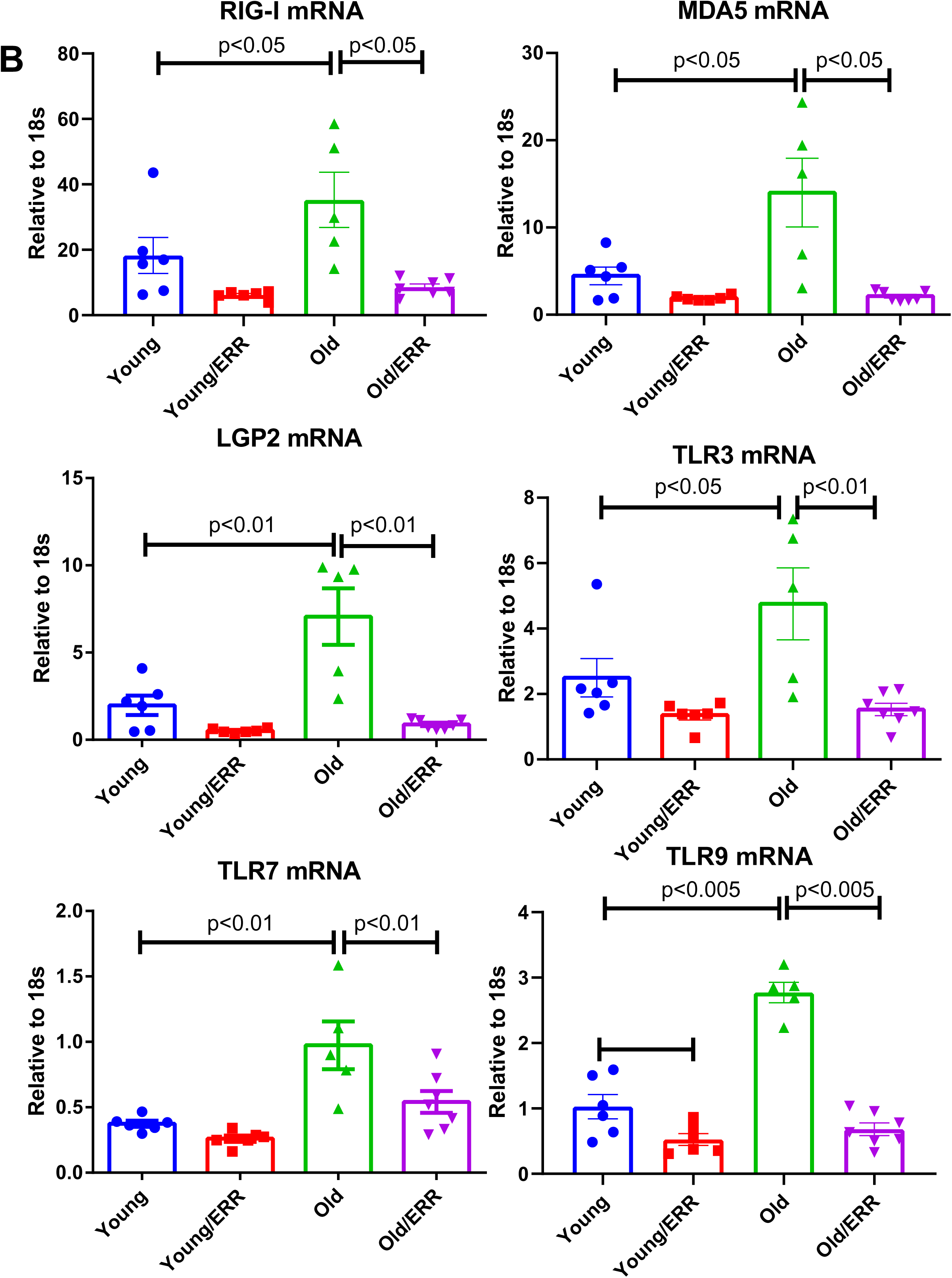
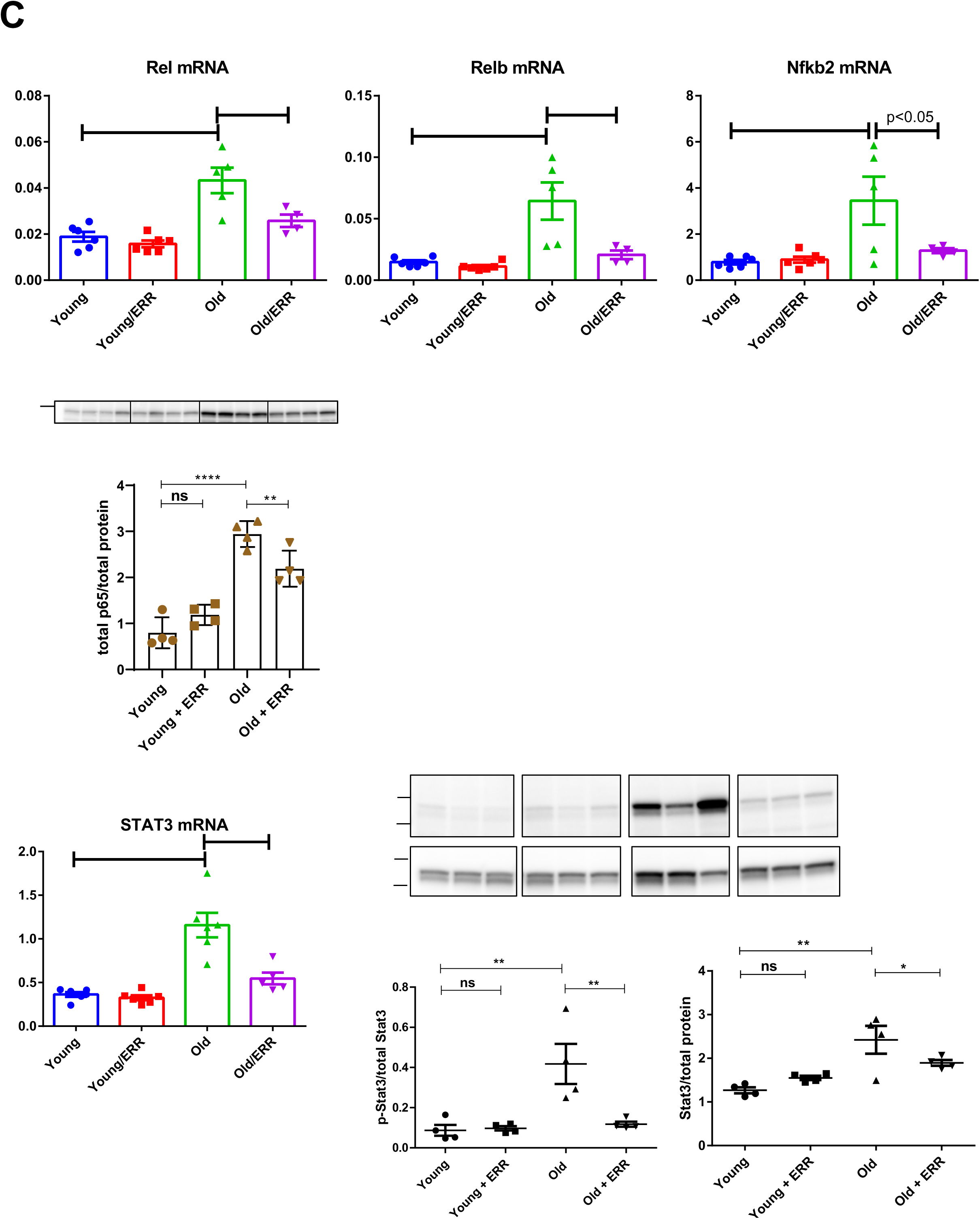
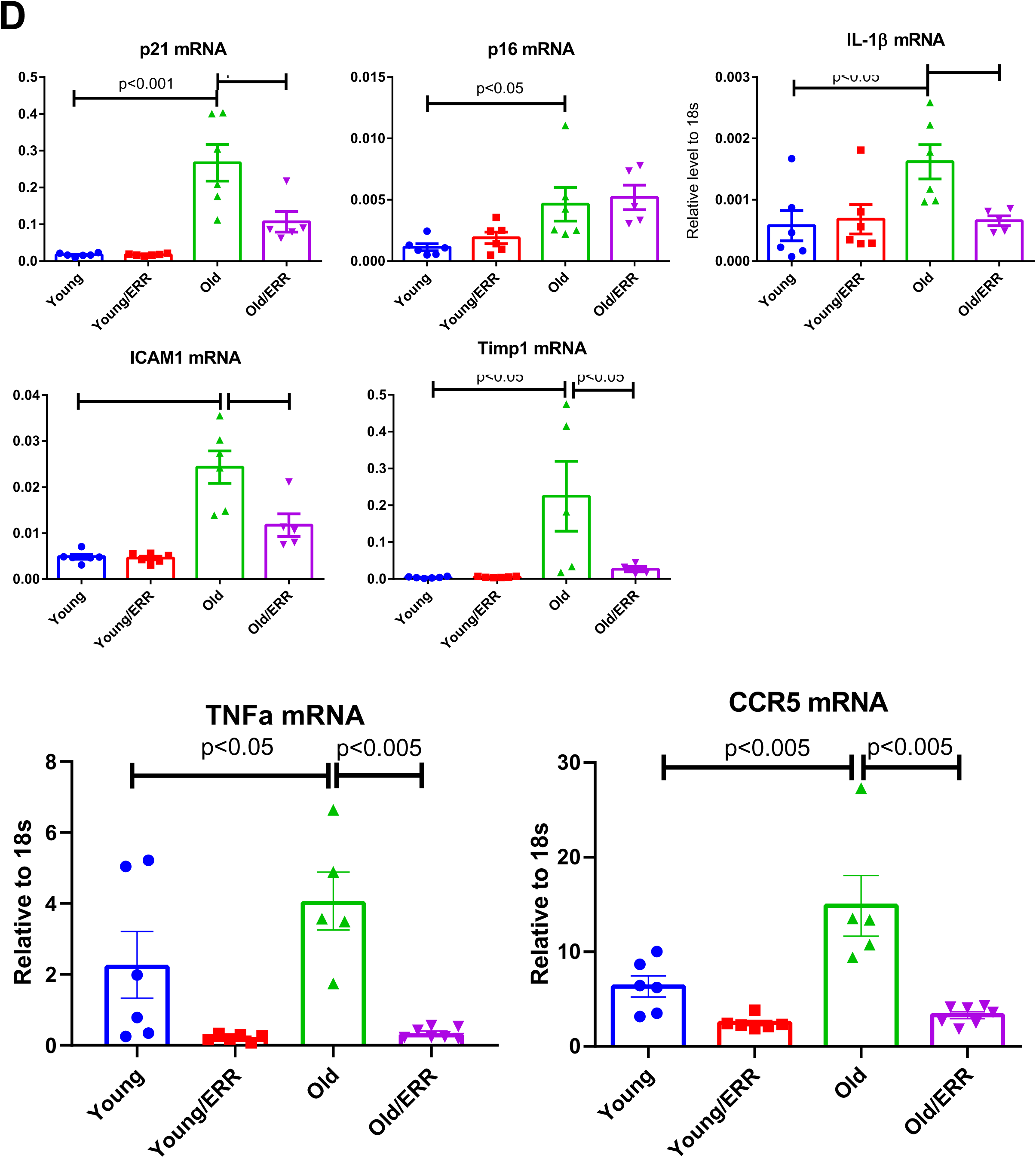
Pan-ERR agonist treatment decreased inflammation in aging kidneys. **A)** There were significant increases in STING and cGAS mRNA and protein level in the kidneys of old mice. Treatment with the pan ERR agonist decreased their expression in the kidneys of old mice. N=5-6 for each group in mRNA level analysis. N=4 for each group in protein analysis. **B)** RIG-I-like receptors (RIG-I, MDA5, LGP2) and Toll-like receptors (TLR3, 7 and 9) mRNA levels were increased in the aging kidney and decreased with the treatment. N=5-6 for each group. **C)** There were significant increases in *Rel, Relb* and *Nfkb2* mRNA, and total p65 protein expression in the kidneys of old mice and treatment with the pan ERR agonist decreased their expression to levels seen in the kidneys of young mice. There were also similar changes to *Stat3* mRNA, and p-Tyr705-STAT3 and total STAT3 protein expression. N=5-6 for each group in mRNA level analysis. N=3-4 for each group in protein analysis. **D)** There were significant increases in the cellular senescence marker p21 and p16 in the aging kidney and the pan agonist treatment reduced the level of p21 expression but not p16. As senescence associated secretory phenotype marker, *Il1b, Icam1, Timp1, Tnfa,* and *Ccr5* mRNA were found increased in the kidneys of old mice and treatment with the pan ERR agonist decreased their expression. N=5-6 for each group.

In addition, we examined another potential STING-activated interferon signaling pathway and observed the corresponding changes in STAT3 (**Figure 7C**). We found significant increases in both total STAT3 and p-Tyr705-STAT3 protein, a marker for STAT3 activation. These increases were substantially suppressed upon treatment with the pan-ERR agonist. This cascade signaling activation was in parallel to the expression of senescence marker p21, which was induced in the aging kidney and reduced by the treatment with the pan-ERR agonist (**Figure 7D**). Another senescence marker, p16, was increased in the aging kidneys but was unaffected by the treatment (**Figure 7D**). Both of these cellular senescence markers are downregulated in aging kidneys with life-long CR (**Supplementary Figure 5**).

To determine which senescence associated secretory phenotype (SASP) factors are regulated by the pan-ERR agonist treatment in the aging kidney, we searched the RNAseq data and verified by real-time PCR that RNAs encoding the proinflammatory cytokines IL-1β and TNF-α, the chemokine receptor CCR5, the metallopeptidase inhibitor TIMP1, and the cell adhesion molecule ICAM1 were increased in the aging kidney and treatment with the pan-ERR agonist decreased their expression (**Figure 7D**).

### STING inhibitor decreased inflammation and increased mitochondrial gene expression in aging kidneys

To determine whether the increase in STING mRNA and protein per se mediates age-related increase in inflammation, we treated aging mice with a known STING inhibitor C-176 ^66^. C-176 decreased IL-1β, Stat3, phospho-Stat3 and the senescence marker p21 expression in the aging kidney (**Figure 8A**). Unexpectedly, we found the expression of master regulator for mitochondria biogenesis PGC1α and PGC1β in aging kidney were also increased with the treatment, along with ERRα (**Figure 8B**). Genes involved in mitochondrial ETC complex, Ndufs1, Cox6a2, and ATP5e, and fatty acid oxidation gene Mcad were also found upregulated by the STING inhibitor (**Figure 8B**).

**Figure 8.**
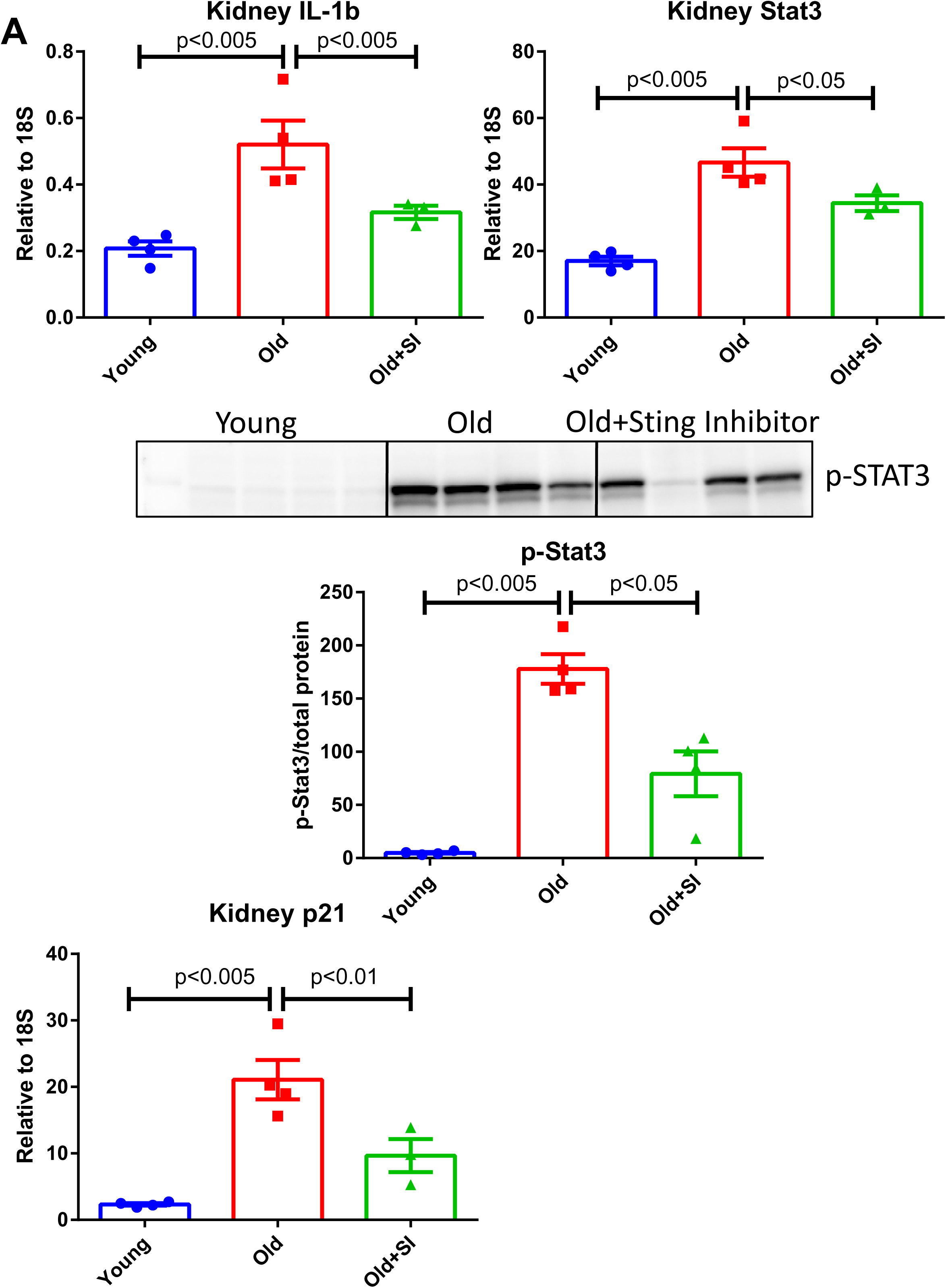
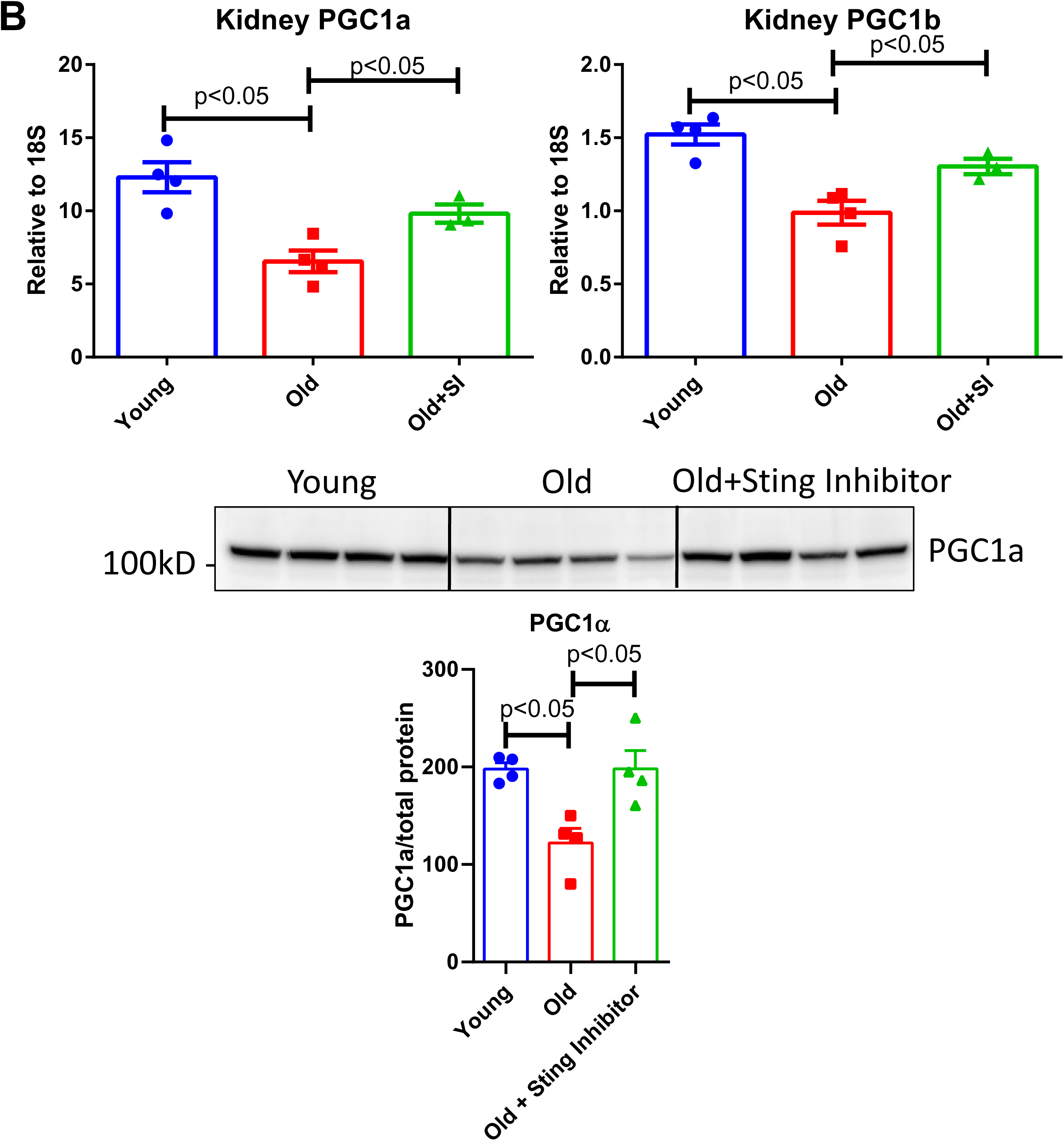
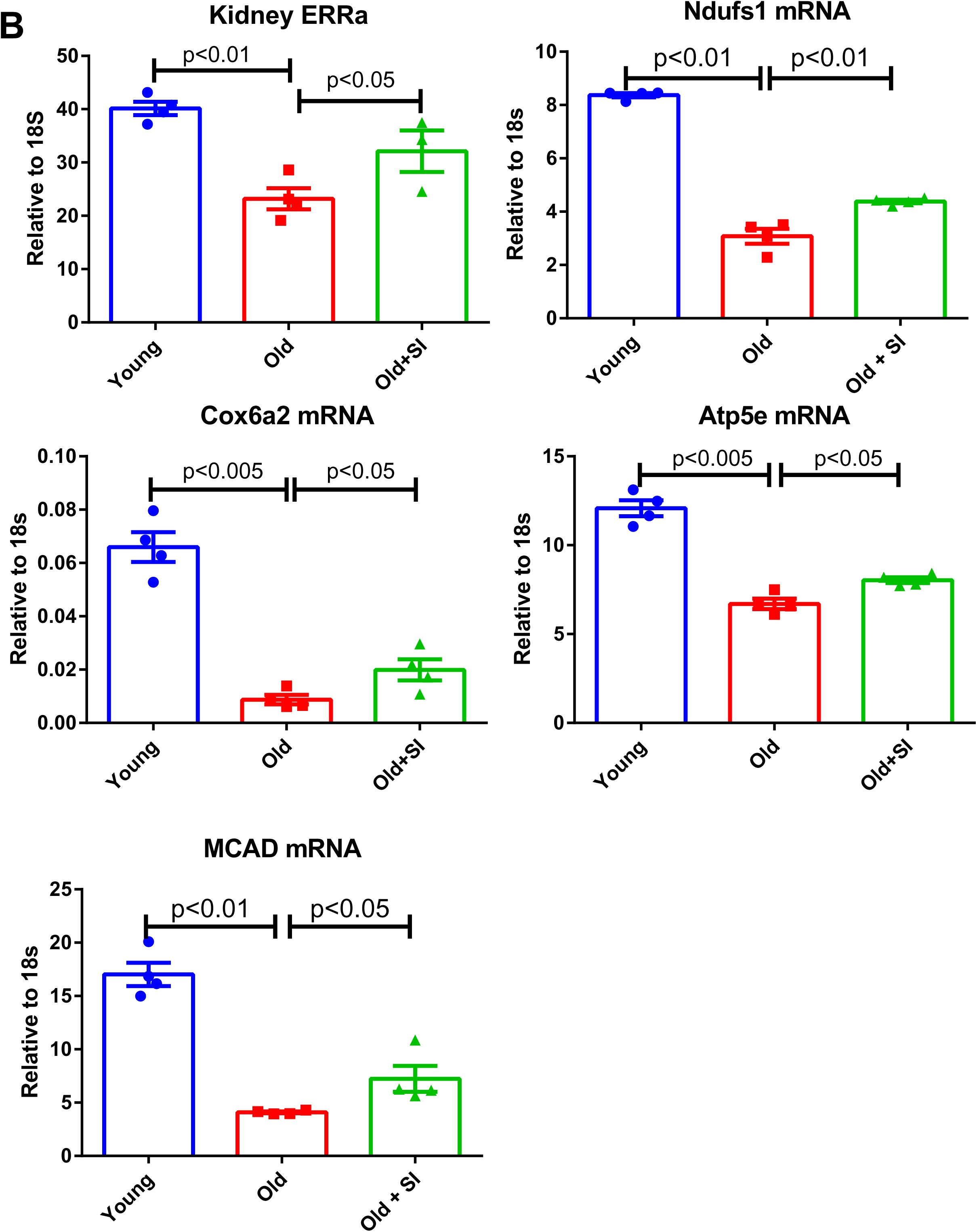
Treatment of aging mice with STING inhibitor C-176. **A)** STING inhibitor decreased IL-1β, Stat3, phospho-Stat3, and p21 expression in aging kidneys. N=4 for each group. **B)** STING inhibitor increased expression of PGC1α, PGC1β, ERRα, Ndufs1(complex I), Cox6a2 (complex IV), Atp5e (complex V), and MCAD in aging kidneys. N=4 for each group.

Interestingly, in proximal tubular epithelial cells, TNF-α, the proinflammatory cytokine which is increased in the aging kidney, induced impairments in genes that mediate mitochondrial biogenesis and the ETC complex (**Figure 1D**). These results suggest that in aging, inflammation and mitochondrial dysfunction may regulate each other to amplify age-related kidney disease.

## DISCUSSION

The data presented here have identified the nuclear hormone receptors the estrogen related receptors ERRα, ERRβ, and ERRγ as important modulators of age-related mitochondrial dysfunction and inflammation. ERRα, ERRβ, and ERRγ expression is decreased in the aging kidney, and lifelong CR results in increases in expression of ERRα, ERRβ, and ERRγ in the kidney. In parallel, CR also prevents age-related mitochondrial dysfunction and inflammation ^12, 67, 68^. ERRs therefore act as potential CR mimetics in the kidney. Remarkably, only an 8-week treatment of 21-month-old mice with the pan-ERR agonist reversed the age-related increases in albuminuria and podocyte loss, mitochondrial dysfunction and inflammation. These effects were comparable with those achieved with lifelong CR, which is known to protect against age-related co-morbidities, including loss of renal function ^10, 11, 67^.

Recent evidence indicates that mitochondrial dysfunction is one of the mediators of cellular senescence, and the associated SASP includes pro-inflammatory cytokines and pro-fibrotic growth factors ^69, 70^. This process may also be involved in the age-related inflammation, termed inflammaging or senoinflammation, which is also prevented by CR ^71, 72^.

Our RNAseq data indicated that apoptosis and cell-cycle regulating pathways were also enriched in the aging kidney. Further analysis of this data in a supervised manner identified 29 senescence associated genes upregulated in the aging kidneys and 10 downregulated. In particular, we found that p53 (Trp53), p65 (Rela), and p21(Cdkn1a) which are the major indicators of senescent cells were highly expressed in aging kidney.

The downregulated genes in aging kidney were enriched in mitochondrial and metabolic processes based on our mRNA sequencing data. Dysregulated mitochondrial functioning has been widely studied in aging kidneys^42^. RNAseq data suggests that Tfam mRNA levels are decreased in aging kidney, which we have verified by RT-QPCR determination. TFAM is a key regulator of mitochondrial gene expression ^45^ and is crucial for maintaining mtDNA structure, transcription, and replication^46^. It has been shown that deletion of Tfam leads to mtDNA escape into the cytoplasm and activation of the innate immune pathway through cGAS-STING activation which plays key roles in immunity, inflammation, senescence and cancer ^44, 63, 73, 74^. In addition to the recent identification of the importance of this signaling pathway in mouse models of acute kidney injury, chronic kidney disease and fibrosis ^43, 44^, our studies also show increased expression of STING in aging kidneys, and its downregulation following treatment with the pan-ERR agonist.

In addition to cGAS-STING signaling, mitochondrial dysfunction is also associated with activation of STAT3 ^75^. Increased STAT3 signaling is associated with senescence ^76^ as well as kidney disease ^77^. We found increased STAT3 in the aging kidneys both at the mRNA and protein level, including increased phospho-STAT3 (Tyr^705^) and normalization after treatment with the pan ERR agonist.

To determine whether STING activation per se mediates aging-related inflammation, we treated aging mice with a STING inhibitor. We found that STING inhibition decreased inflammation in aging kidney, as reported in other models using a STING inhibitor or STING knockout mice ^43, 44^. However, we found that STING inhibition can further increase mitochondrial gene expression, suggesting a model where mitochondrial injury triggers inflammation, compounding mitochondrial dysfunction in aging kidney. Interestingly, in proximal tubular epithelial cells, TNF-α, the proinflammatory cytokine which is increased in the aging kidney, induced impairments in genes that mediate mitochondrial biogenesis and the ETC complex. These results suggest that in aging, inflammation and mitochondrial dysfunction may regulate each other to amplify age-related kidney disease. Our findings are consistent with reports of mitochondrial damage in acute kidney injury induced by lipopolysaccharide (LPS), an endotoxin that activate innate immunity (via TLR4) to induce circulating cytokines ^78–80^.

Overall, we found that in aging kidney there is a dysregulation of mitochondrial and immune processes. Pan-ERR agonist treatment mitigated several of the abovementioned pathways. In summary, our studies identify the ERRs as beneficial modulators of mitochondrial dysfunction and inflammation in the aging kidney.

## ACKNOWLEDGEMENTS

The above study is supported by NIA R01 AG049493 and NIDDK R01 DK116567 to M.L. and AHA postdoctoral fellowship to K.M. (19POST34381041) and A.E. L. (19POST34430001). J.B.K. is supported by the NIDDK Intramural Research Program. F.J.G and U.G. are supported by the NCI Intramural Research Program.

## AUTHOR CONTRIBUTIONS

This study was conceived and led by M.L. and X.X.W. X.X.W. performed most of the laboratory work. K.M. assisted with the animal studies and biochemical analysis. Y.Q. and U.G. performed the proteomics and assisted with the analysis. J.P. A.T., V.S., B.O., and L.B. performed the multi-omics processing and integration analyses. S.R. and K.B. conducted the FLIM imaging and analysis. P.D. and A.Z.R. performed the immunohistochemistry and analysis. A.E.L. performed human primary cell culture work. P.Z. and J.B.K. performed the EM and B.A.J., K.B. and A.Z.R. assisted with the data analysis. T.B. provided the reagent pan-ERR agonist. X.X.W. and M.L. wrote the manuscript with editorial input from all authors.

## DATA AVAILABILITY STATEMENT

The data that support the findings of this study are available from the corresponding author upon reasonable request. RNAseq raw data have been deposited in NCBI GEO database with accession number: PRJNA642362. Proteomics data are available via ProteomeXchange with identifier PXD020051.

## Supplementary Materials Table of Contents

Supplementary Methods ……………………………………………………………………………………………………………….. 2

Supplementary Figures …………………………………………………………………………………………………………………. 6

Supplementary Table 1 ………………………………………………………………………………………………………………… 16

For supplementary tables 2-7 please refer to additional supporting documents (Supplementary tables 2-7.rar).

