## supplementary methods, figures and table 1 for "Estrogen-related receptor agonism reverses mitochondrial dysfunction and inflammation in the aging kidney"

### Supplementary Material Table of Contents

For supplementary tables 2-7 please refer to additional supporting documents (Supplementary tables 2-7.rar).

#### SUPPLEMENTARY METHODS

**Autofluorescence FLIM measurements using DIVER microscope**: To determine the effects of ERR activation on aging kidney, autofluorescence images from kidney slices of 5 µm thickness were measured using an Olympus microscope equipped with a DIVER (Deep Imaging via Enhanced Recovery) detector developed at the Laboratory for Fluorescence Dynamics (LFD), University of California at Irvine, CA. This microscope has a higher photon collection efficiency due to the large area detector and the photon counting nature of it enables FLIM imaging. The details of the detector construction have been described previously (1-6). The system assembled in Georgetown University is based on an Olympus FVMPERS (Olympus, Waltham, MA) upright laser scanning microscope equipped with the special DIVER detector. A short-pulsed tunable ultrafast (fs) laser (Insight Deep See X3, Spectra-Physics, Santa Clara, CA) is used as the excitation source. The samples were excited with a 20X 0.45NA air objective (Olympus, Waltham, MA). The sample is placed directly on top of the detector assembly input window below the objective and two-photon induced fluorescence is detected by a large area photomultiplier (PMT) (Hamamatsu R7600P-300, Campbell, CA). The detector assembly consists of a sealed chamber with the filter wheel/shutter inside and the housing with PMT. The refractive index matching liquid filled inside the housing removes loss of photons due to internal reflections and thus achieves efficient collection of photons. Two BG-39 filters serve as input and output windows of the chamber, and block NIR excitation light from entering PMT, transmitting UV and visible fluorescence and harmonic signals. The only optical elements in the detector assembly are BG-39 filters and the glass filter of the filter wheel, allowing detection of emitted photons from 320-650 nm wavelength range. The samples were excited with a 710 nm pulsed laser and two photon autofluorescence FLIM was collected using the blue filter of the detection assembly ($\lambda_{EM}$ = 410 -510 nm). Each of the images was taken with a 625 µm field of view, 20 µs pixel dwell time and 16 repeat scans for increasing signal to noise ratio.

The signal from the PMT is collected using a FLIMBox (ISS, Champaign, IL) and directly transferred to the phasor plot (Figure S1). Details of the phasor approach towards FLIM analysis has been explained elsewhere (7-10). Briefly, this method of lifetime analysis involves transferring the fluorescence intensity decays in the Fourier space and plotting the Fourier components against each other. This results in a phasor plot (Figure S1) where each point results from a pixel of the image. The autofluorescence from the tissue have a non-zero lifetime and appear inside the semicircle. A distribution in the phasor plot can then either be selected using a continuous color scale and the images can be colored accordingly. Phasor approach towards FLIM is a fit free approach, increases the speed of the analysis, and decreases the computational difficulty associated with FLIM technique (7-11). The data collection and analysis was carried out by using SimFCS, developed by Prof. Enrico Gratton at the LFD (https://www.lfd.uci.edu/globals/). A continuous color scheme was used for the phasor-mapped image and the redder/ purple color represents more bound NADH and more cyan/white color represents more free NADH (Figure 5E in main text). The position of the bound NADH and free NADH are shown by red and blue cursors, respectively (Figure S1) (12).

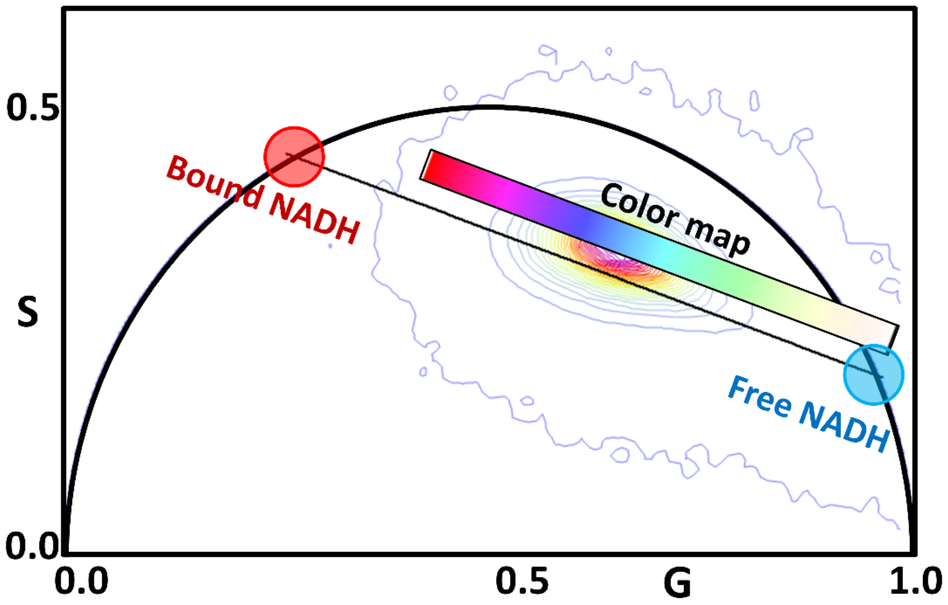

**Figure S1:** The phasor map showing the lifetime distribution of NADH in kidney slices. A continuous color scheme was used for the phasor-mapped image and the redder/ purple color represents more bound NADH and more cyan/white color represents more free NADH. The position of the bound NADH and free NADH are shown by red and blue cursors, respectively.

### Supplementary Figure 1

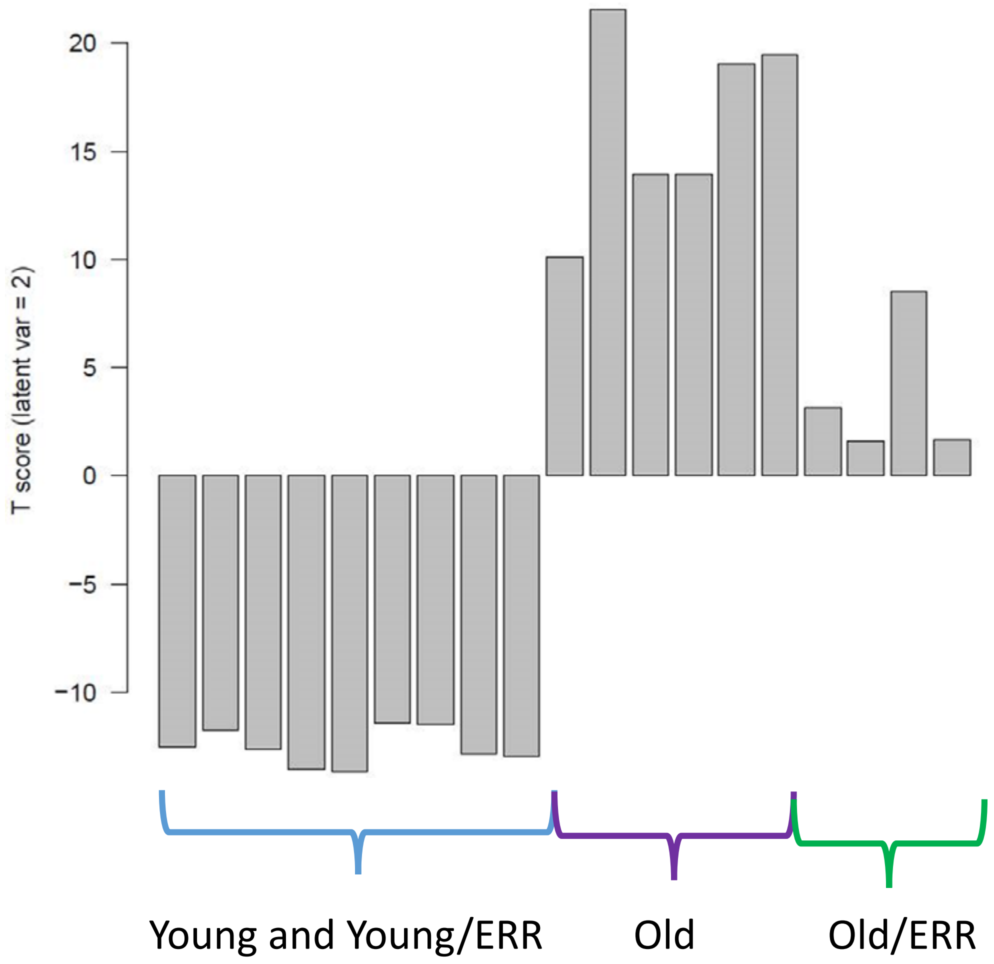

**Supplementary Figure 1:** O2PLS component (V2) profile separates the samples into three biologically meaningful groups: kidneys of young mice treated with vehicle or with pan-ERR agonist, kidneys of old mice, and kidneys of old mice treated with pan-ERR agonist.

### Supplementary Figure 2

A.

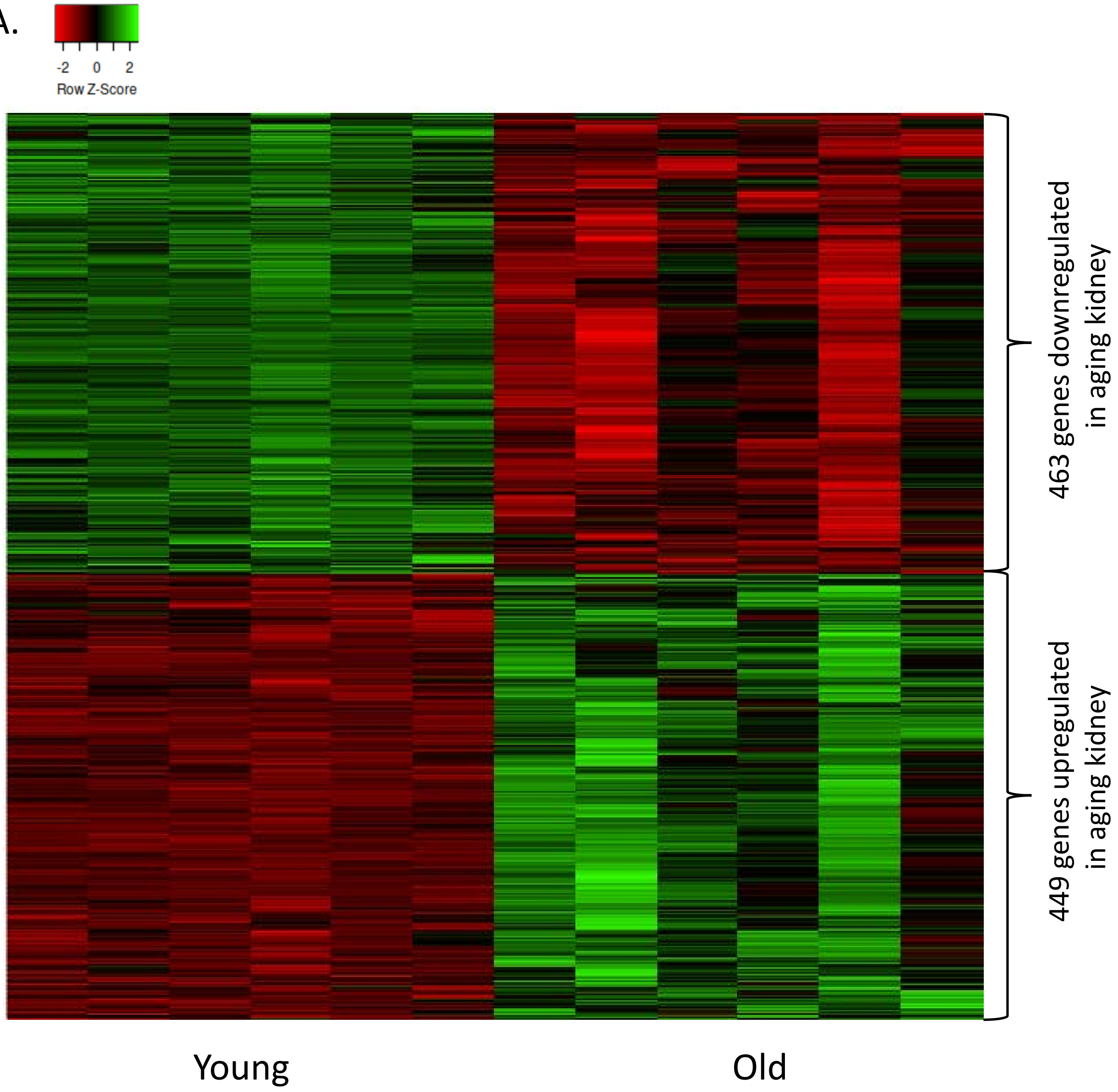

**Supplementary Figure 2:** RNA-seq and proteomics analysis of kidneys from old mice compared with kidneys from young mice

**A)** Heat map showing expression patterns of genes differentially expressed in kidneys of old mice compared to kidneys of young mice. The heat map indicates up-regulation (green), down-regulation (red),and unaltered gene expression (black). The columns represent individual samples.

B.

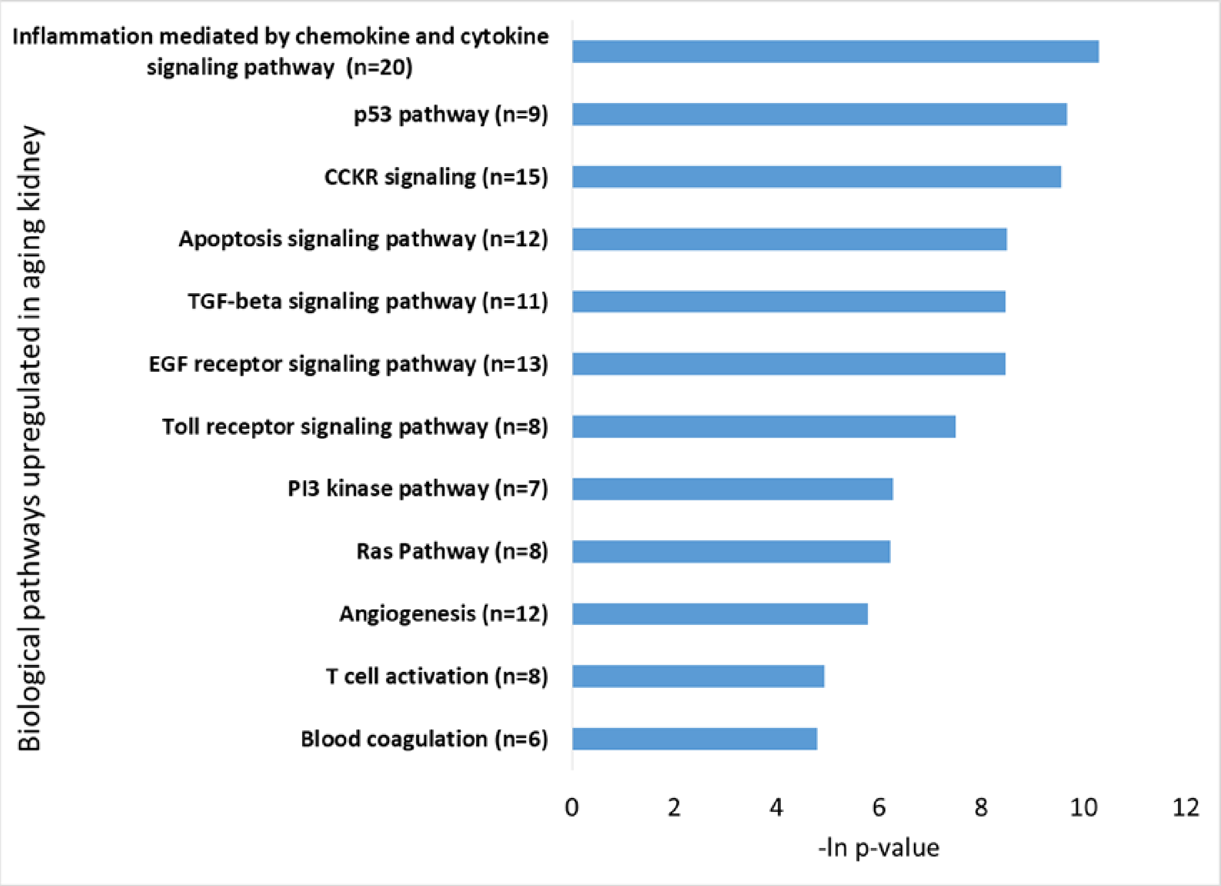

**Supplementary Figure 2: B)** Functional pathway enrichment analysis of genes upregulated in kidneys of old mice compared to kidneys of young mice. The y-axis shows significantly enriched pathways. The x-axis indicates p-value of enrichment of the given pathway.

C.

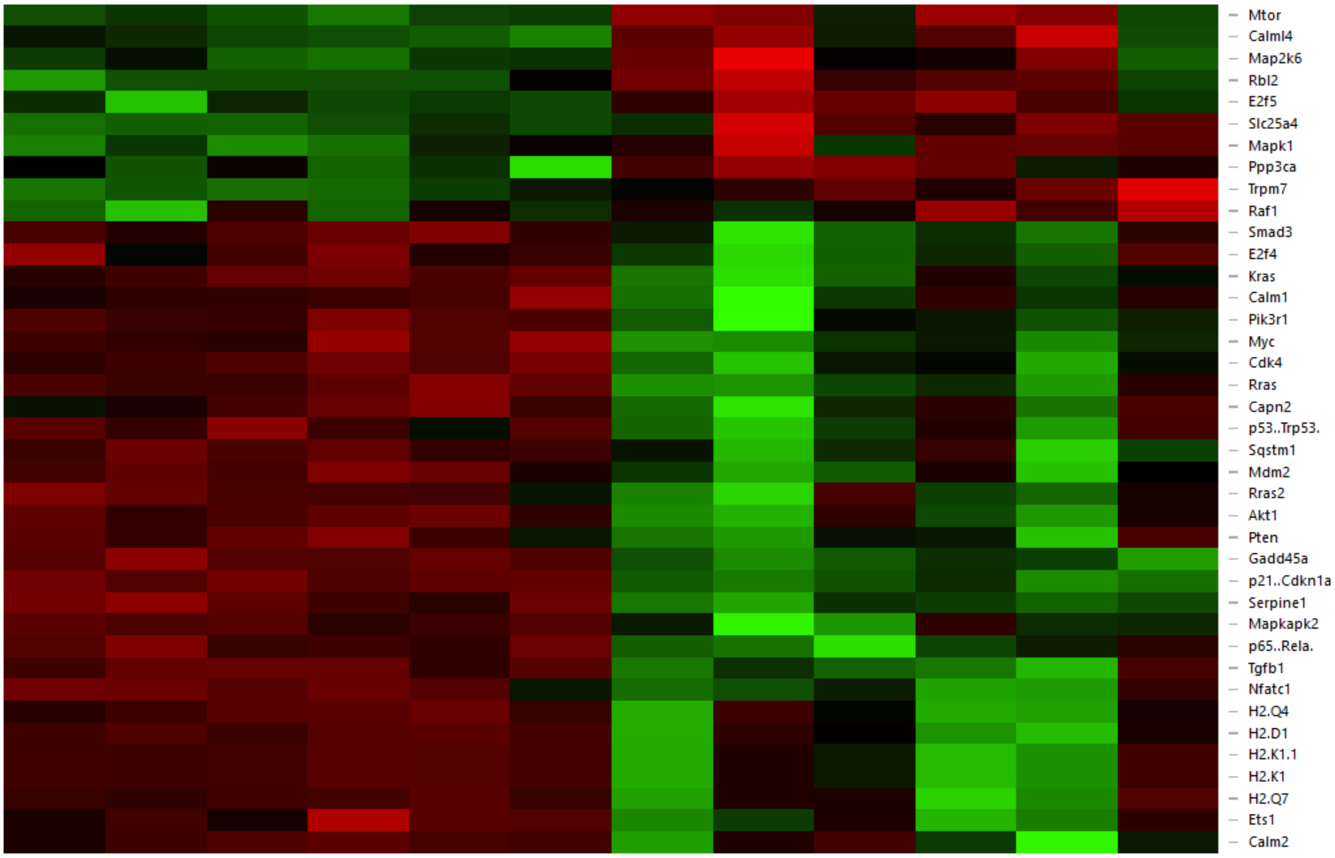

**Supplementary Figure 2: C)** Heat map showing expression patterns of Senescence related genes differentially expressed in kidneys of old mice compared to kidneys of young mice. The heat map indicates up-regulation (green), down-regulation (red), and unaltered gene expression (black). The columns represent individual samples.

D.

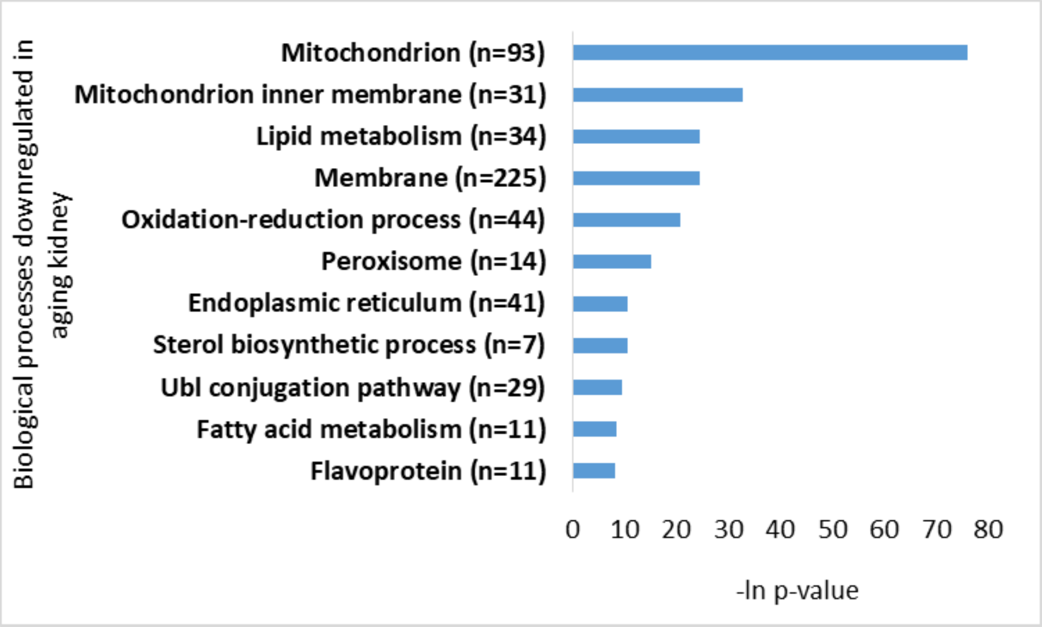

E.

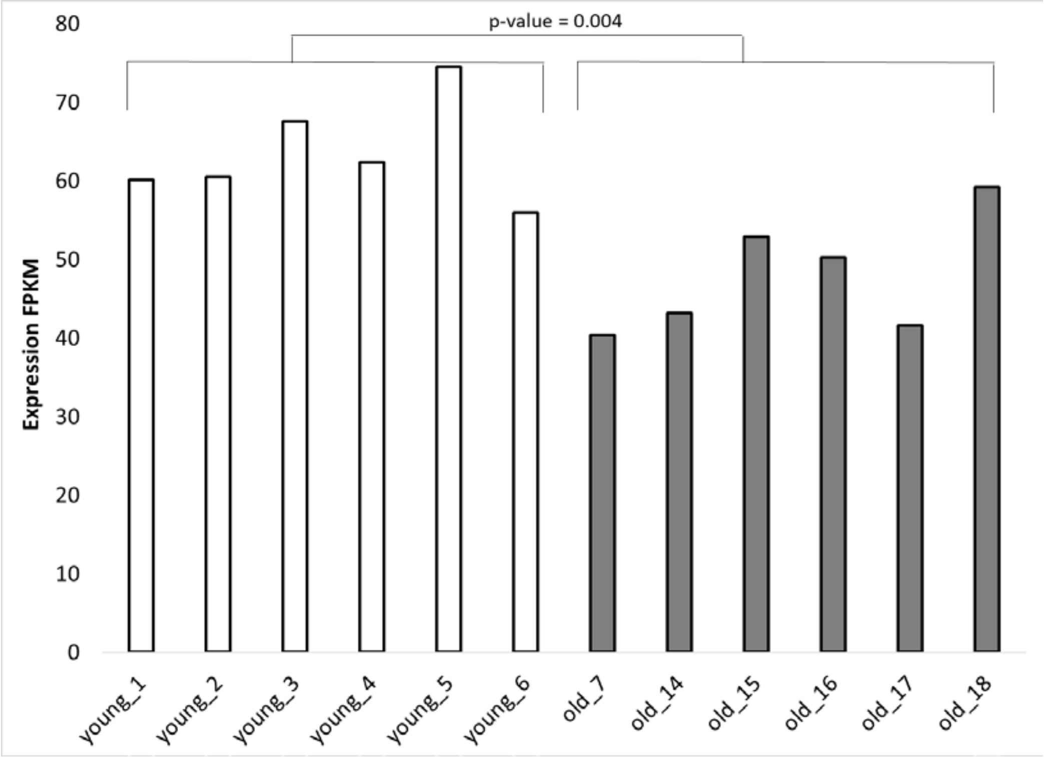

**Supplementary Figure 2: D)** Functional pathway enrichment analysis of genes downregulated in kidneys of old mice compared to kidneys of young mice. The y-axis shows significantly enriched pathways. The x-axis indicates p-value of enrichment of the given pathway. **E)** Tfam mRNA expression profile in young and old kidneys. Expression values are presented in FPKM units.

### Supplementary Figure 2

F.

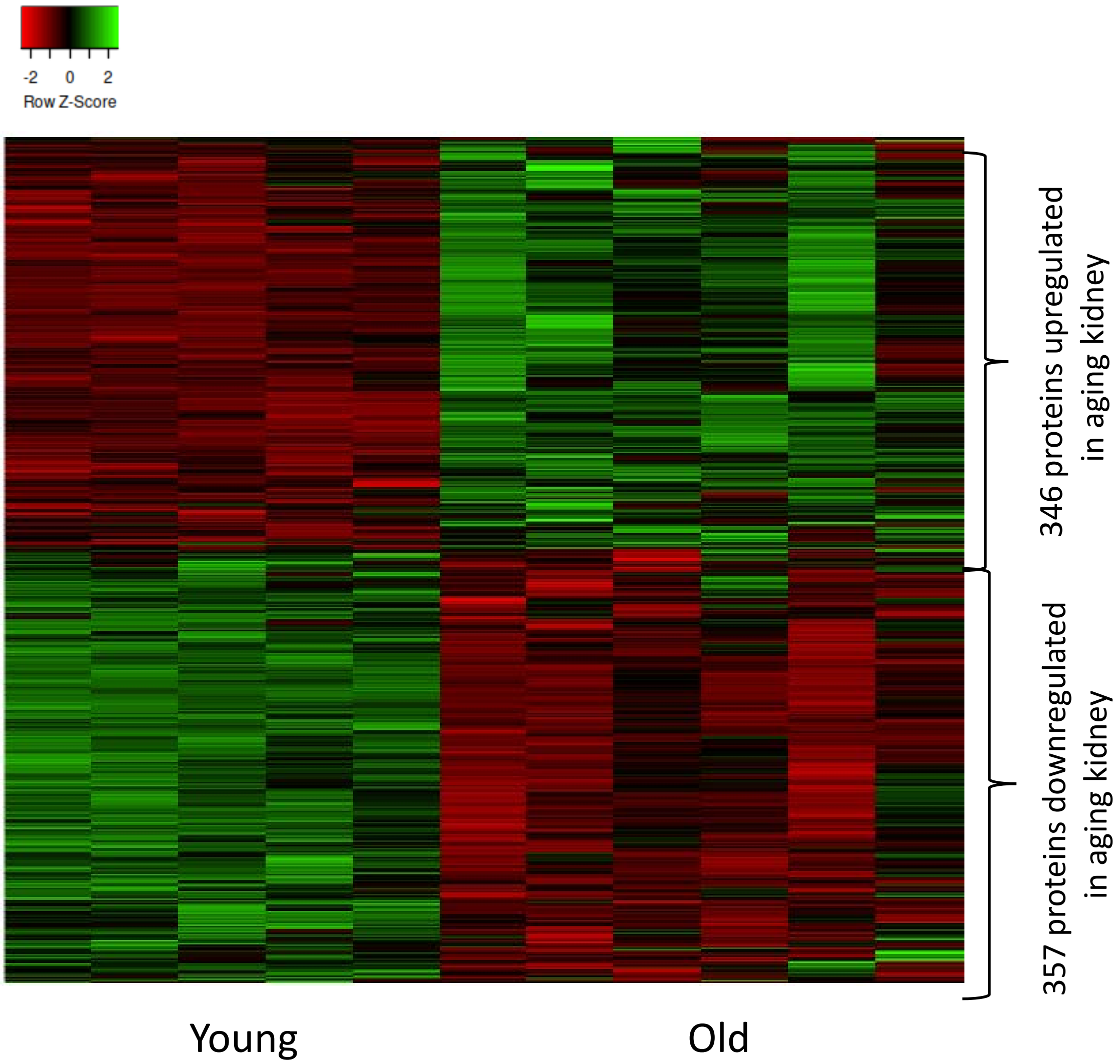

**Supplementary Figure 2: F)** Heat map showing expression patterns of proteins differentially expressed in kidneys of old mice compared to kidneys of young mice. The heat map indicates up-regulation (green), down-regulation (red), and unaltered gene expression (black). The columns represent individual samples.

G.

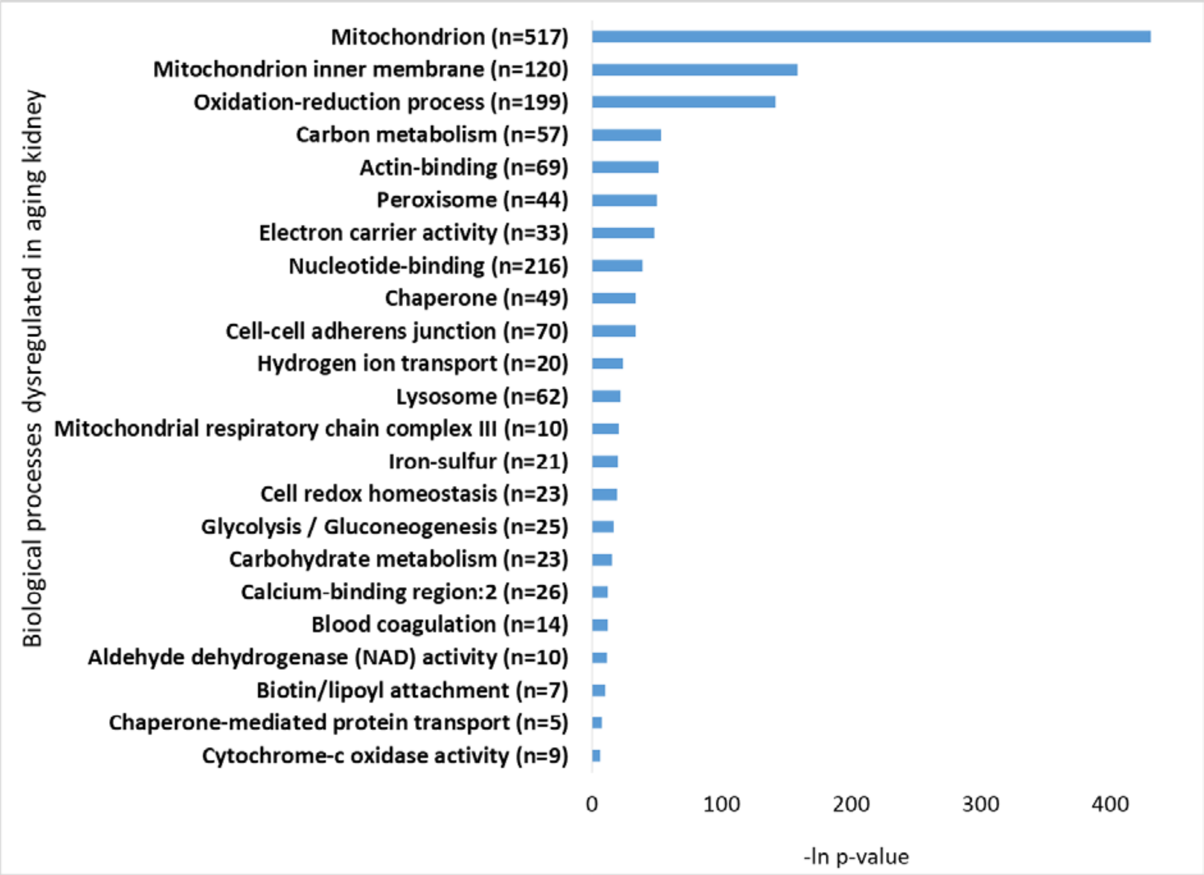

**Supplementary Figure 2: G)**Functional pathway enrichment analysis of differentially expressed proteins in kidneys of old mice compared to kidneys of young mice. The y-axis shows significantly enriched pathways. The x-axis indicates p-value of enrichment of the given pathway.

### Supplementary Figure 3

**PCA: proteome abundance (pool-ratio quantile normalized) before batch correction**

**PCA: proteome abundance (pool-ratio quantile normalized) after batch correction**

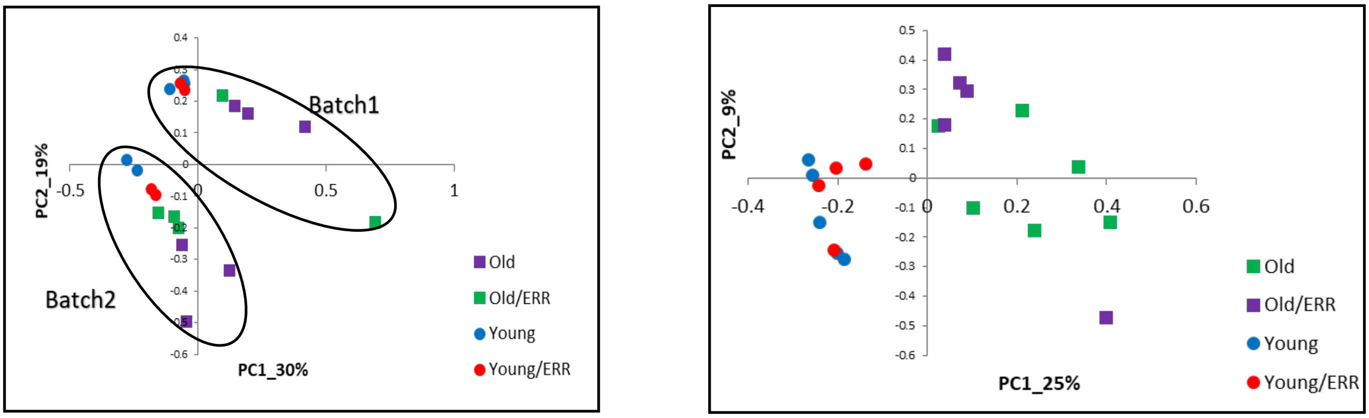

**Supplementary Figure 3:** Principle component analysis of proteomics data prior to batch correction procedure and after implementing Experimental Bayes (EB) batch correction method. After batch correction, the samples on the PC1-PC2 plane are separated into biologically meaningful groups.

### Supplementary Figure 4

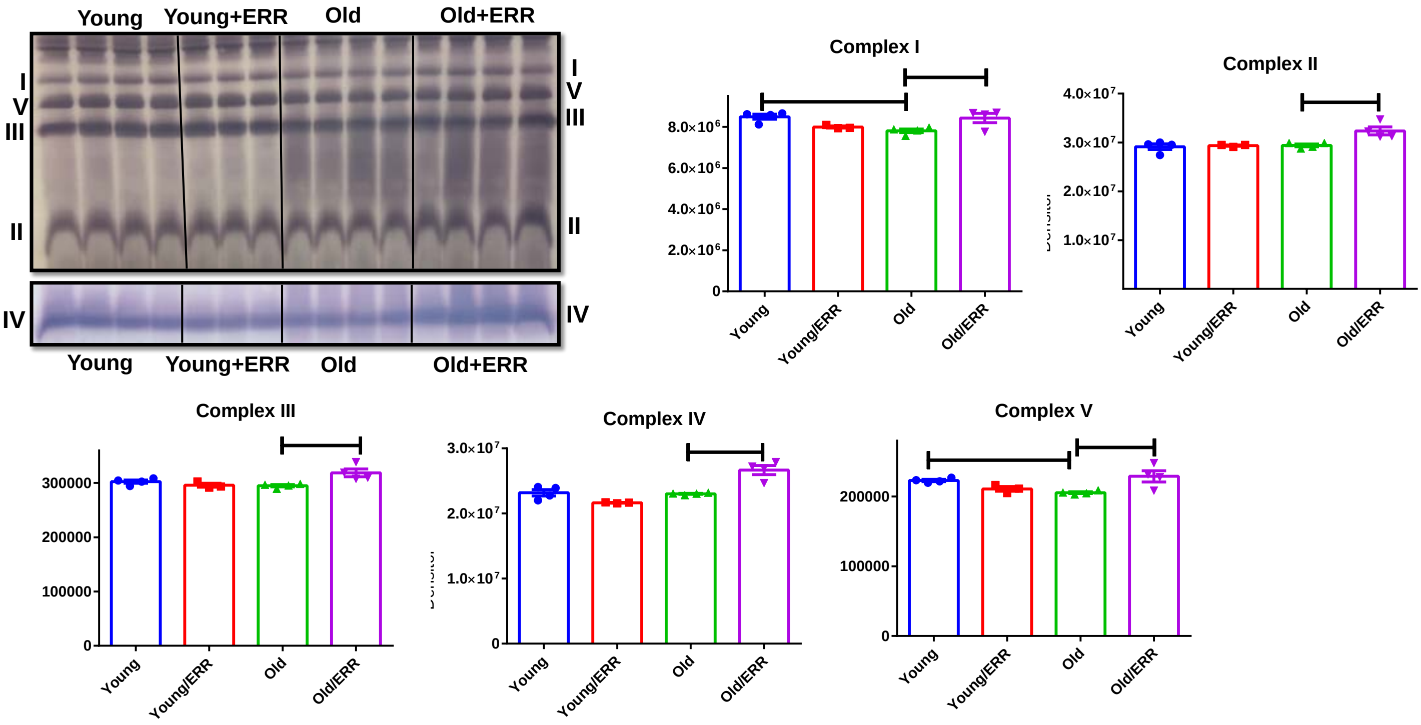

**Supplementary Figure 4:** Native blue gel indicates the increased level of assembled complex I, II, III, IV and V in the kidneys of old mice after treatment with the pan ERR agonist. N=4 for each group.

### Supplementary Figure 5

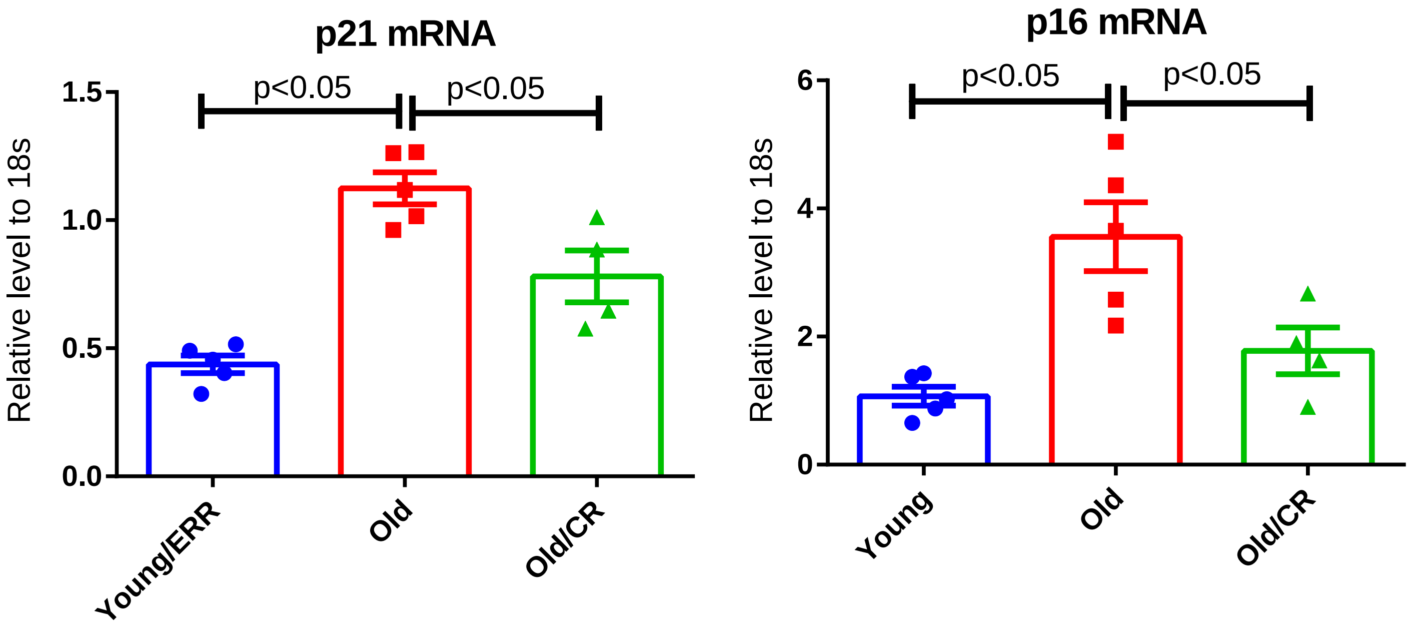

**Supplementary Figure 5:** Cellular senescence markers p21 and p16 mRNA level was increased in the aging kidney. However in aging kidneys with life-long CR, p21 and p16 mRNA expression was downregulated compared to that in *ad lib* aging kidneys. n=4-5 samples per group.

**Supplementary Table 1:**

**A. List of primers**

| Name | Forward | Reverse |
|---|---|---|
| *Esrrα* | CAGGGAGGGAAGGGATGG | ATGAGGAGAGGAGCGAAGG |
| *Esrrb* | GCACCTGGGCTCTAGTTGC | TACAGTCCTCGTAGCTCTTGC |
| *Esrrg* | AAGATCGACACATTGATTCCAGC | CATGGTTGAACTGTAACTCCCAC |
| *Ppargc1a* | GTCAGAGTGGATTGGAGTTG | AAGTCATTCACATCAAGTTCAG |
| *Ppargc1b* | TCCTGTAAAAGCCCGGAGTAT | GCTCTGGTAGGGGCAGTGA |
| *Pdk4* | AGGGAGGTCGAGCTGTTCTC | GGAGTGTTCACTAAGCGGTCA |
| *Acadm* | AACACTTACTATGCCTCGATTGCA | CCATAGCCTCCGAAAATCTGAA |
| *Tfam1* | AACACCCAGATGCAAAACTTTCA | GACTTGGAGTTAGCTGCTCTTT |
| *mtDNA* | ATAACCGAGTCGTTCTGCCAAT | TTTCAGAGCATTGGCCATAGAA |
| *Sdhc* | GCTGCGTTCTTGCTGAGACA | ATCTCCTCCTTAGCTGTGGTT |
| *Atp5b* | GGTTCATCCTGCCAGAGACTA | AATCCCTCATCGAACTGGACG |
| *Pdhb* | AGGAGGGAATTGAATGTGAGGT | ACTGGCTTCTATGGCTTCGAT |
| *Mdh1* | TTCTGGACGGTGTCCTGATG | TTTCACATTGGCTTTCAGTAGGT |
| *Idh3b* | TGGAGAGGTCTCGGAACATCT | AGCCTTGAACACTTCCTTGAC |
| *Sucla2* | ACCCTTTCGCTGCATGAATAC | CCTGTGCCTTTATCACAACATCC |
| *Cpt1a* | CTCCGCCTGAGCCATGAAG | CACCAGTGATGATGCCATTCT |
| *Ndufb8* | TGTTGCCGGGGTCATATCCTA | AGCATCGGGTAGTCGCCATA |
| *Opa1* | CGACTTTGCCGAGGATAGCTT | CGTTGTGAACACACTGCTCTTG |
| *Miga2* | GGAGGACTGAGGGTATGTCCA | CAAGGGCTGTGGCAAAAAGA |
| *Pld6* | ACCTGCACCGAGGCTTTAC | CATGTAGTCGCAGTCAGTGATG |
| *Il1b* | GCAACTGTTCCTGAACTCAACT | ATCTTTTGGGGTCCGTCAACT |
| *Icam1* | GTGATGCTCAGGTATCCATCCA | CACAGTTCTCAAAGCACAGCG |
| *Stat3* | AGCTGGACACACGCTACCT | AGGAATCGGCTATATTGCTGGT |
| *Uqcrb* | GGCCGATCTGCTGTTTCAG | CATCTCGCATTAACCCCAGTT |
| *Cox6a2* | CTGCTCCCTTAACTGCTGGAT | GATTGTGGAAAAGCGTGTGGT |
| *Tmem173* | GGTCACCGCTCCAAATATGTAG | CAGTAGTCCAAGTTCGTGCGA |
| *Cdkn1a* | CCTGGTGATGTCCGACCTG | CCATGAGCGCATCGCAATC |

**B. Antibodies**

| Name | Host | Source | Catalogue Number |
|---|---|---|---|
| OPA1 | Mouse | BD Biosciences | 612606 |
| MFN2 | Rabbit | Millipore | 978-715-4321 |
| DRP1 | Mouse | Novusbio | H00010059-M01 |
| p-DRP1 (s616) | Rabbit | Cell Signaling | 3455S |
| Stat3 | Rabbit | Cell Signaling | 4904T |
| p-Stat3 (Y705) | Rabbit | Cell Signaling | 9145T |
| PDH e2/e3 | Mouse | Abcam | Ab110333 |

For **supplementary tables 2-6** please refer to additional supporting documents (Supplementary tables 2-6.rar).
